# Differential expression and homotypic enrichment of a classic Cadherin directs tissue-level contractile asymmetry during neural tube closure

**DOI:** 10.1101/425165

**Authors:** Hidehiko Hashimoto, Edwin Munro

## Abstract

Embryos pattern force generation at tissue boundaries during morphogenesis, but how they do so remains poorly understood. Here we show how tissue-specific expression of the type II cadherin, Cadherin2 (hereafter Cad2), patterns actomyosin contractility along the neural/epidermal (Ne/Epi) boundary to drive zippering and neural tube closure in the basal chordate, *Ciona robusta*. Cad2 is differentially expressed and homotypically enriched in neural cells along the Ne/Epi boundary, where RhoA and Myosin are activated during zipper progression. Equalizing Cad2 expression across the Ne/Epi boundary inhibits RhoA/Myosin activation and zipper progression, while creating ectopic Cad2 expression boundaries is sufficient to direct RhoA/Myosin activity to those boundaries. We show that Cad2 polarizes RhoA activity by sequestering the Rho GTPase activating protein, Gap21/23, to homotypic junctions, which in turn redirects RhoA/Myosin activity to heterotypic Ne/Epi junctions. By activating Myosin II along Ne/Epi junctions ahead of zipper and inhibiting Myosin II at new Ne/Ne junctions behind zipper, Cad2 promotes tissue level contractile asymmetry to drive zipper progression.

## Introduction

Actomyosin contractility has emerged as a key force generator for cell movement and shape change in embryonic development. Contractile forces produced locally by individual cells are transmitted through cell-cell adhesions, and resolved more globally as tissue deformation and remodeling (reviewed in (Dahmann et al., 2011; Fagotto, 2015; Heisenberg and Bellaiche, 2013; Lecuit et al., 2011)). Thus, a fundamental challenge is to understand how contractility is patterned in space and time to orchestrate diverse tissue-level morphogenetic movements such as invagination, elongation, and dynamic maintenance of tissue boundaries (reviewed in (Fagotto, 2015; Heisenberg and Bellaiche, 2013)).

Work to date suggests that embryos pattern contractility by controlling local activation of myosin II in space and time, using two general types of mechanism. The first involves local control of myosin II with respect to intrinsic apico-basal or planar polarity within a particular tissue or compartment. For example, apical localization and/or activation of factors such as Fog, T48 and Shroom family members can promote local activation of Myosin II and apical constriction to drive tissue bending/invagination (reviewed in (Gilmour et al., 2017; Heisenberg and Bellaiche, 2013; Lecuit et al., 2011; Martin and Goldstein, 2014)). Alternatively, polarized activation of Myosin II at cell-cell junctions by core members of the Planar Cell Polarity (PCP) pathway can drive planar-polarized junction contraction, leading to cell-cell intercalation and tissue elongation (reviewed in(Harris, 2018; Heisenberg and Bellaiche, 2013; Shindo, 2018; Walck-Shannon and Hardin, 2014)). For example, in chick embryos, local enrichment of PCP proteins Celser1, on mediolaterally oriented apical junctions, promotes planar polarized myosin activation, junction contraction and cell intercalation to induce tissue invagination and elongation during neural tube closure (Nishimura et al., 2012).

A second class of mechanisms involves the differential expression of signaling or cell-adhesion molecules across tissue or compartment boundaries. The majority of examples thus far come from studies in Drosophila. For example, differential expression of the nectin-like homophilic binding protein Echinoid induces the formation of actomyosin cables at tissue boundaries during dorsal appendage formation in egg chambers and dorsal closure in embryos (Laplante et al., 2006; Laplante and Nilson, 2011). Similarly, local expression of the homophilic binding protein Crumbs within the salivary gland placode promotes the assembly of an actomyosin cable at the placode boundary to help drive invagination (Röper, 2012). Differential expression of Toll family receptors in stripes along the AP axis bias myosin activity to mediolaterally oriented junctions to promote cell intercalation and germband elongation (Paré et al., 2014). Finally, differential expression of signaling molecules such as Wingless, Hedgehog, Dpp and Notch across compartment boundaries in embryos and imaginal discs, activates Myosin II to prevent cell mixing across those boundaries (reviewed in (Dahmann et al., 2011)). Recent studies in chordates highlight a key role for differential expression of Eph/Ephrin signaling across multiple tissue boundaries in localized activation of myosin II and the dynamic maintenance of tissue separation ((Fagotto et al., 2013), reviewed in (Fagotto, 2015)). While a number of other molecules have been implicated in the control of contractility at tissue boundaries, their contributions remain poorly defined (Fagotto, 2015). More generally, how differential expression of signaling molecules across tissue boundaries leads to locally polarized activation of myosin II within single cells remains poorly understood.

Neural tube closure in ascidians offers a powerful opportunity to study the dynamic control of Myosin II along tissue boundaries during epithelial morphogenesis. Neurulation is one of the defining events of chordate morphogenesis, in which the neural tube forms and separates from surface epidermis to form the rudiment of the future nervous system. In ascidians, the rudiment includes an axial nerve cord and a central vesicle, corresponding to the spinal cord and brain respectively, of higher vertebrates (Lemaire et al., 2002; Schoenwolf and Smith, 1990; Wallingford et al., 2013; Yamaguchi and Miura, 2012); light blue and dark blue in Figure 1A). Like many vertebrates, ascidians form a neural tube in three steps: (1) the neural plate invaginates, raising neural folds along its lateral boundary with the epidermis (the Ne/Epi boundary; Figure 1A); (2) convergent extension brings the neural folds closer to the presumptive midline (Navarrete and Michael Levine, 2016) and (3) local meeting of the neural folds, and local rearrangements of apical junctions (Ne/Epi -> Ne/Ne plus Epi/Epi), separate a closed neural tube from a continuous overlying epidermis (Hashimoto et al., 2015; Nicol and Meinertzhagen, 1988a; 1988b; Ogura and Sasakura, 2016). This final step proceeds directionally from posterior to anterior, and has thus been referred to as zippering (Hashimoto et al., 2015; Nicol and Meinertzhagen, 1988b; 1988a); Figure 1A.

**Figure 1.**
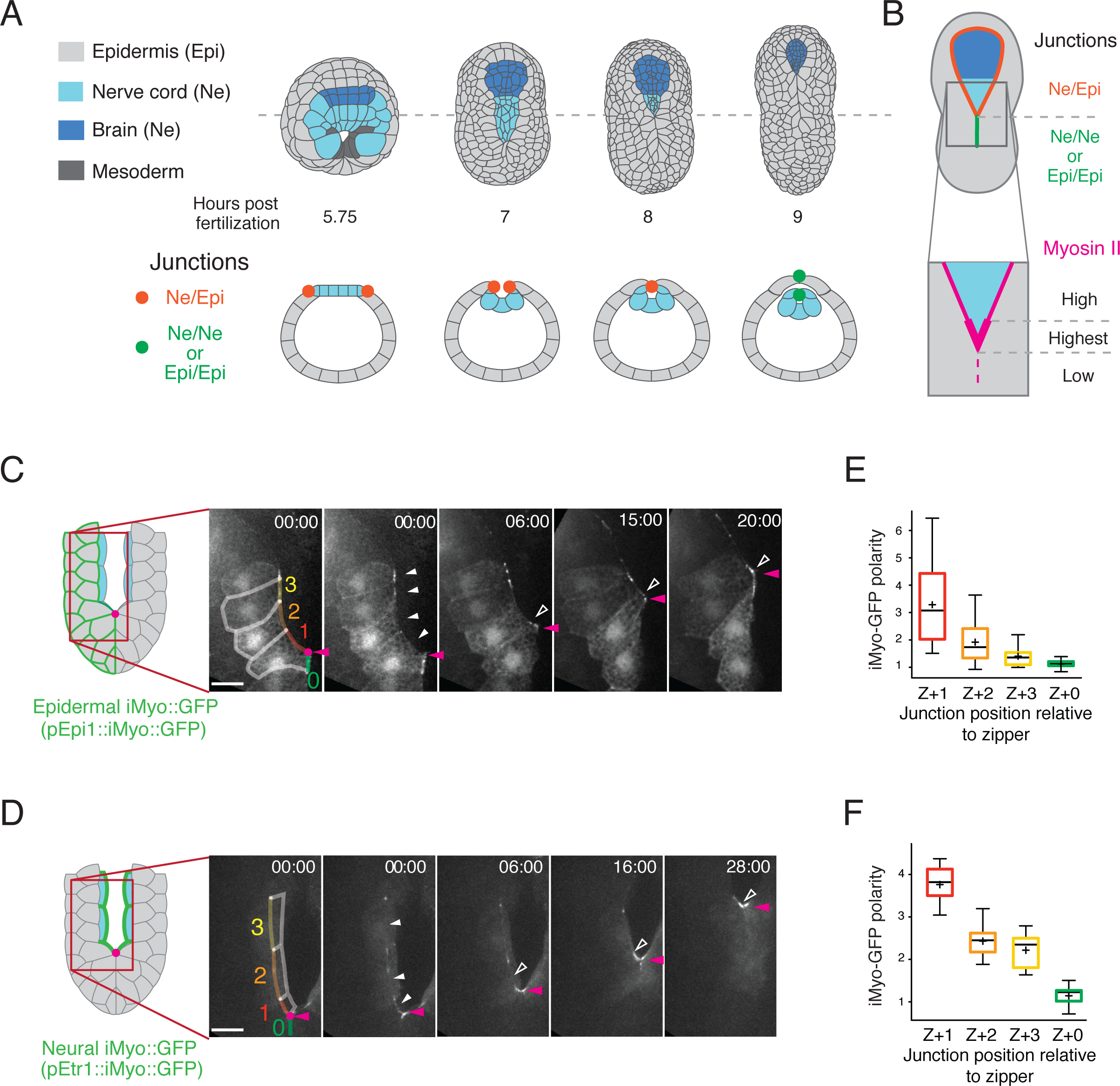
Dynamic accumulation of myosin along the Ne/Epi boundary during zippering. (A) Schematic overview of neural tube closure. Top row shows dorsal view and bottom row shows cross-sectional view of embryos at 5.75, 7, 8 and 9 hours after fertilization. Dashed line in top row indicates the axial position of cross sections in bottom row. Anterior is up. (B) Top: heterotypic Ne/Epi boundaries (orange) ahead of the zipper; homotypic Ne/Ne and Epi/Epi junctions (green) behind the zipper. Bottom: distribution of myosin activity (magenta) measured previously by antibody staining: low on homotypic boundaries behind the zipper, high on heterotypic boundaries ahead of the zipper, and highest just ahead of the zipper. (C-F) Spatiotemporal patterns of myosin accumulation in neural and epidermal cells. (C, D) Frames from Movies S1 showing distribution of iMyo⸬GFP when expressed under the control of epidermal-specific and neural-specific promoters. Schematics at left indicate the domains of expression. Magenta arrowheads indicate zipper position. White arrowheads indicate increased myosin along the entire Ne/Epi boundary (filled arrows) and just ahead of the zipper (open arrows). Note that expression of iMyo⸬GFP is mosaic and restricted to the left half of the embryo in C, and to nerve cord cells flanking Ne/Epi junctions in D (see also Figure S1C and S1D). Scale bars = 5μm. (E, F) Quantitation of average iMyo⸬GFP polarity in epidermal (E) and neural (F) cells as a function of cell position relative to zipper, indicated by color overlays in the first movie frames in C and D. Box plots show the distribution of average polarity values measured for individual junctions during the period in which an Ne/Epi junction next to the zipper (z + 1) starts and finishes its contraction. Horizontal line indicates median, + indicates mean, boxes indicate second and third quartiles, and whiskers indicate 95% confidence interval. (n = 23 junctions, from 8 embryos in E and n = 12, from 6 embryos in F; see Materials and Methods for details).

Previously, we showed that spatiotemporal control of myosin activity along the Ne/Epi boundary plays a key role in driving zippering and neural tube closure (Hashimoto et al., 2015). Active myosin is enriched on all Ne/Epi junctions ahead of the zipper, and it is reduced on newly-formed Ne/Ne or Epi/Epi junctions behind the zipper (Figure 1A and 1B). In addition, strong sequential activation of Myosin II on Ne/Epi junctions just ahead of the zipper drives a posterior to anterior sequence of rapid junction contractions (Figure 1B). Computer simulations suggest that this tissue-level asymmetry in myosin activity (high ahead of the zipper and low behind the zipper) is essential to ensure that sequential contractions are converted efficiently into forward movement of the zipper (Hashimoto et al., 2015).

Here, we identify a molecular basis for this tissue-level asymmetry. We show that a classical Cadherin2 (hereafter Cad2, Figure S1A) is expressed specifically in neural cells, where it localizes preferentially to homotypic Ne/Ne junctions. Cad2, in turn, recruits the putative Rho GAP (Gap21/23, orthologue of human ARHGAP21 and 23) to Ne/Ne junctions and away from Ne/Epi junctions to pattern local activation of RhoA and Myosin II. By activating Myosin II along the Ne/Epi boundary ahead of the zipper, and inhibiting Myosin II at newly-formed Ne/Ne contacts behind the zipper, the Cad2/Gap21/23/RhoA pathway defines a tissue-level feedback loop, supporting a tissue-level contractile asymmetry that dynamically adjusts itself to a moving zipper to coordinate zippering and neural tube closure. We suggest that dynamic coupling of junction exchange to local changes in contractility may play key roles in controlling fusion and separation of epithelia in many other contexts.

## Results

### Myosin accumulates along the Ne/Epi boundary in both Neural and Epidermal cells

We previously showed that Myosin II accumulates along the entire Ne/Epi boundary during zippering (Hashimoto et al., 2015), but we did not determine in which cells (Ne or Epi or both) this accumulation occurs. To address this question as a first step towards identifying upstream control mechanisms, we expressed a GFP-tagged intrabody (“iMyo-GFP”), that recognizes the non-muscle Myosin II heavy chain (Hashimoto et al., 2015; Nizak et al., 2003) under the control of either neural-specific promoter pEtr1 ((Veeman et al., 2010) Figure S1A and S1B) or epidermal-specific promoter pEpi1 ((Sasakura et al., 2009), Figure S1A and S1B). Then we used live imaging to examine dynamic localization of iMyo-GFP during zipper progression. We found that in both neural and epidermal cells, iMyo-GFP was enriched along the entire Ne/Epi boundary (white fill arrowheads in Figure 1C and 1D; Movie S1), and highly enriched on rapidly contracting junctions just ahead of the advancing zipper (white open arrowheads in Figure 1C and 1D; quantified in Figure 1E and 1F, S1C and S1D). Thus, myosin activity is patterned in both neural and epidermal cells during zipper progression.

### Cad2 is differentially expressed and homotypically enriched in midline neural cells

To identify mechanisms that polarize myosin activity along Ne/Epi boundary, we searched for genes encoding putative transmembrane proteins that are differentially expressed across the Ne/Epi boundary during zippering. We focused on the midline cells – two rows of neural and epidermal cells that flank the Ne/Epi boundary. The midline cells arise from a single pair of bilaterally symmetric founder cells called b6.5. The b6.5 cells cleave at the 64-cell stage to produce daughters b7.9 and b7.10 (Figure 2Ai), which divide unequally during gastrulation to produce pairs of neural (b8.17 and b8.19) and epidermal (b8.18 and b8.20) precursor cells (Figure 2Aii). The neural and epidermal precursor cells divide once and twice more, respectively, along the AP axis, to produce two rows of neural and epidermal midline cells (Figure 2Aiii).

**Figure 2.**
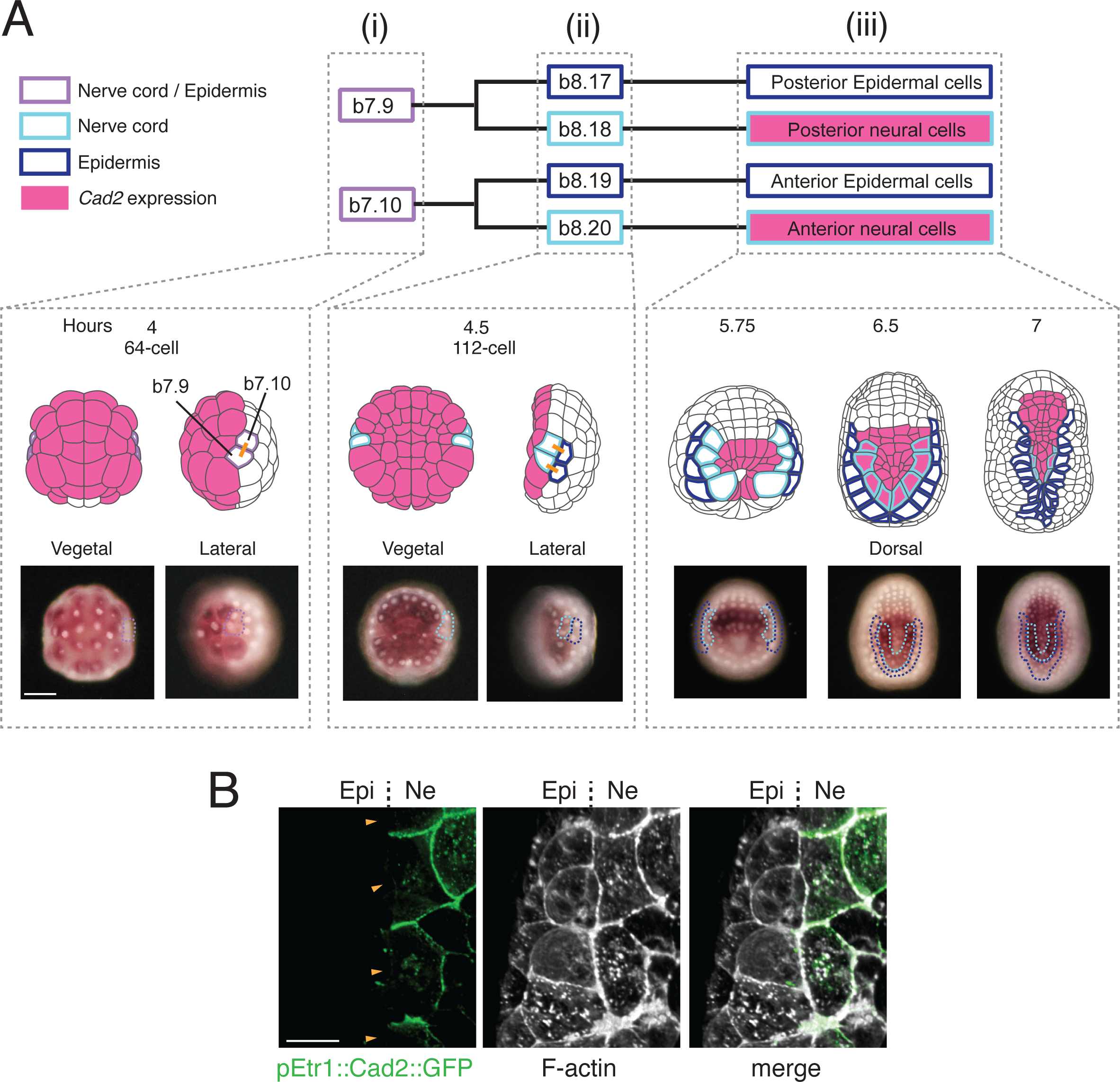
Stage and lineage-specific expression of *Cad2*. (A) Lineage tree for midline nerve cord and epidermal cells. Midline Ne/Epi precursors (purple) divide at ~4 hours after fertilization to produce founders of the midline neural (light blue) and epidermal (dark blue) lineages. Magenta fill indicates cells expressing *Cad2*. Top: Schematic views showing the positions of cells at the indicated stages. Orange bars indicate the orientation of a previous cleavage. Bottom: Distribution of *Cad2* mRNA at the same stages. Midline neural/epidermal precursors and their progeny are outlined in color, as in (A). Anterior is up in all panels. Scale bars, 50 μm. (B) Zoomed view of neural cells ahead of the zipper expressing Cad2⸬GFP under the control of a neural-specific Etr1 promoter. Embryos were fixed and counter-stained with phalloidin. Orange arrowheads indicate the Ne/Epi boundary. Scale bars = 5 μm.

We searched the ascidian database ANISEED for genes that are expressed before the onset of zippering in the descendents of either b8.17 and b8.19, or b8.18 and b8.20. This search identified Cadherin2 (Cad2, also known as *Cadherin.b*, Figure S1A), one of two classical cadherins in the *Ciona robusta* genome (Sasakura et al., 2003). *Cad2* encodes a Type II Cadherin most closely related to Human VE-Cadherin (CHD5) (Hulpiau and van Roy, 2009) and Cadherin-8 (CHD8) (Sasakura et al., 2003), which was previously shown to be expressed in the neural primordium (Noda and Satoh, 2008). To better characterize the spatiotemporal pattern of *Cad2* expression, we performed simultaneous in situ hybridization and nuclear staining. Before gastrulation, *Cad2* is expressed throughout the vegetal hemisphere in most neural precursor cells, but not in the neural midline precursors b7.9 and b7.10 ((Noda and Satoh, 2008); Figure 2Ai, ii). *Cad2* is first expressed in neural midline cells b9.33, b9.34, b9.37 and b9.38 just before initiation of zippering (Figure 2Aiii), suggesting a specific role in neural tube closure. Because commercially available antibodies do not recognize endogenous Cad2, we expressed a GFP-tagged form of Cad2 using the neural-specific promoter pEtr1, and examined its subcellular localization. We found that Cad2⸬GFP was enriched at homotypic junctions between Cad2⸬GFP-expressing neural cells ahead of the zipper (Figure 2B), and newly-formed Ne/Ne junctions behind the zipper (Figure S6A). In contrast, Cad2⸬GFP was absent from heterotypic junctions between Cad2⸬GFP-expressing neural cells and non-expressing epidermal cells (Figure 2B; quantified in S6A). We refer to this pattern of enrichment, in which Cad2 is enriched at contacts between Cad2-expressing cells, but not at contacts between expressing and non-expressing cells, as homotypic enrichment.

### Differential expression and homotypic enrichment of Cad2 directs myosin activity to expression boundaries

If differential expression and homotypic enrichment of Cad2 in midline neural cells directs myosin activity to heterotypic Ne/Epi junctions (Figure 3A), then equalizing *Cad2* expression across the Ne/Epi boundary should abolish homotypic enrichment and reduce or abolish myosin activity, while creating ectopic *Cad2* expression boundaries in neural or epidermal territory should induce homotypic enrichment and activate myosin at those boundaries. We tested these predictions in two ways. First, we used a midline-specific promoter, pMsx (Roure et al., 2014), to express Cad2⸬GFP in all midline cells (Figure S1B), forcing similarly high levels of Cad2 expression across the Ne/Epi boundary and creating an ectopic expression boundary in epidermal territory (Figure 3B). Second, we used lineage-specific injection of morpholinos to inhibit Cad2 expression exclusively in midline neural cells, forcing similarly low levels of Cad2 expression across the Ne/Epi boundary, and creating an ectopic expression boundary between midline and non-midline neural cells (Figure 3C).

**Figure 3.**
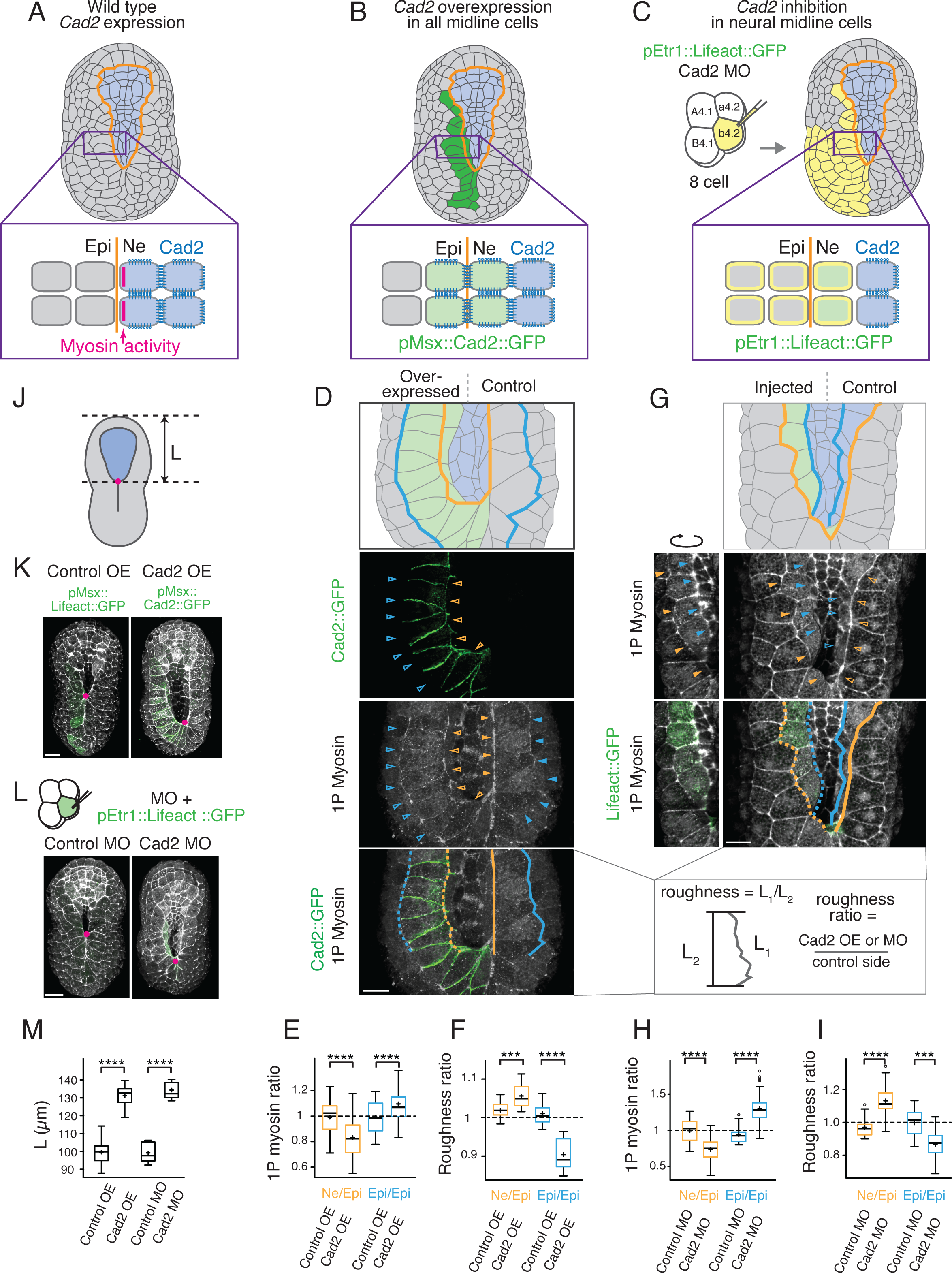
Myosin is activated at *Cad2* expression boundaries. (A) Schematic view of a wild type embryo near the onset of zippering indicating the Ne/Epi boundary (orange line) and *Cad2* expression pattern (blue). Expanded view shows wild type distributions of *Cad2* and myosin activity in neural cells. (B) Cad2⸬GFP (green) is over-expressed in all midline cells on one side of the embryo. (C) Co-injection of Morpholino Oligonucleotides (MO) against Cad2 with an expression marker (pEtr1⸬Lifeact⸬GFP) into a single b4.2 blastomere at the 8-cell stage. Yellow indicates b4.2 descendants; green indicates neural b4.2 descendants that received the MO. (D-F) Effects of ectopic Cad2⸬GFP expression. (D) Top panel shows schematic dorsal view of a neurula over-expressing Cad2⸬GFP (green) over endogenous Cad2 (blue) in midline cells. Color fill indicates ectopic Cad2⸬GFP (green) over endogenous Cad2 (blue). Orange line: original Ne/Epi boundary. Blue line: boundary between midline and non-midline epidermal cells. Bottom three panels show localization of Cad2⸬GFP, 1P-myosin, and their superposition, respectively, in the same embryo. Open and filled arrowheads: corresponding boundaries on Cad2⸬GFP expressing and non-expressing sides of the embryo. Dashed and solid lines indicate the boundary sections measured for roughness, as shown at the right. Scale bars = 5 μm. (E-F) Box plots showing ratios of (E) 1P myosin and (F) ratio of boundary roughness on Cad2⸬GFP (or Lifeact⸬GFP control) expressing:non-expressing sides of the embryo for the indicated boundary types (n > 9 embryos for each condition in E,F). (G-I) Effects of Cad2 knockdown in midline neural cells. (G) Top panel: schematic dorsal surface view of a neurula stage embryo, injected as shown in (C), showing neural cells inheriting co-injected morpholino and GFP marker (green) over endogenous Cad2 (blue). Orange lines: original Ne/Epi boundary. Blue lines: ectopic boundary between midline and non-midline neural cells. Second panel: localization of 1P-myosin in the same embryo. Open and filled arrowheads indicate corresponding boundaries on injected and non-injected sides of the embryo. Third panel: superposition of GFP marker and 1P-myosin. Dashed and solid lines indicate the boundary sections measured for roughness, as shown below. Scale bars = 5 μm (H-I) Box pots showing ratios of (H) 1P myosin and (I) boundary roughness on control MO or Cad2 MO-injected:non-injected sides of the embryo for the indicated junction or boundary types. (n > 9 embryos for each condition). (J-M) Measurements of zippering defects in Cad2⸬GFP over-expressing or MO-injected embryos. (J) Schematic showing measurement of “un-zippered” length L. (K-L) Representative phalloidin-stained embryos showing effects of Cad2⸬GFP over-expression (K) or Cad2 MO injection (L). Control embryos on left; perturbed embryos on right. Scale bars = 10 μm. (M) Quantification of zipper progression, measured as “unzippered length” L (n > 9 embryos for each condition) **p < 0.05, *** p < 0.005, **** p < 0.0005, Student’s t test.

#### Overexpression of Cad2 in midline cells inhibits myosin activation on Ne/Epi junctions, zippering and neural tube closure

When over-expressed in all midline cells, Cad2⸬GFP was enriched on junctions between midline neural and epidermal cells (open orange arrowheads in Figure 3D), and it was absent from junctions between expressing and non-expressing epidermal cells (open blue arrowheads in Figure 3D). Thus over-expressing Cad2 in all midline cells abolishes homotypic enrichment in midline neural cells and induces homotypic enrichment in midline epidermal cells, consistent with the idea that homophilic engagement determines homotypic enrichment.

To determine how the loss or gain of homotypic enrichment affects myosin activity we examined the distribution of Ser19-phosphorylated Myosin II (1P Myosin) in fixed immunostained embryos over-expressing Cad2⸬GFP in the midline cells. We exploited mosaic expression of the electroporated transgenes (Zeller et al., 2006) to make direct comparisons between cells overexpressing Cad2⸬GFP with paired controls on the opposite sides of the same embryo (Figure 3B,D). Strikingly, we found that 1P Myosin was sharply reduced on junctions between neural and epidermal cells over-expressing Cad2⸬GFP, and 1P Myosin was increased on junctions between Cad2⸬GFP-expressing and non-expressing epidermal cells, relative to paired controls (Figure 3D; quantified in 3E). Thus, equalizing *Cad2* expression across the Ne/Epi boundary abolishes homotypic enrichment of Cad2 in midline neural cells, and reduces myosin activity along the Ne/Epi boundary. In contrast, creating an ectopic Cad2⸬GFP expression boundary in the epidermis induces homotypic enrichment of Cad2⸬GFP, and promotes activation of myosin II along the expression boundary (Figure 3D; quantified in 3E).

To assess how the changes in myosin activity described above affect junction tension, we measured boundary “roughness”, defined as the total length of a stretch of boundary divided by the straight-line distance between its endpoints (bottom panel in Figure 3D). Reduced boundary roughness correlates with higher boundary tension in other embryos (Aliee et al., 2012; Landsberg et al., 2009). In embryos over-expressing Cad2⸬GFP in midline cells, we observed increased boundary roughness (indicating reduced tension) along the Ne/Epi boundary where 1P myosin is reduced, and decreased boundary roughness (indicating increased tension) along boundaries between Cad2 expressing and non-expressing epidermal cells, where 1P myosin is increased (bottom right panel in Figure 3D; quantified in Figure 3F). Significantly, these changes in boundary tension were accompanied by a complete loss of zipper progression and neural tube closure (Figure 3J, 3K and 3M). Thus equalizing Cad2 expression across the Ne/Epi boundary by over-expression in midline cells is sufficient to block myosin activation and decreased tension along the Ne/Epi boundary, and to block zipper progression and neural tube closure.

#### Inhibition of Cad2 in midline neural cells inhibits myosin activation on Ne/Epi junctions, zippering and neural tube closure

In a second complementary set of experiments, we injected antisense morpholino oligonucleotides directed against Cad2 (Cad2 MOs) at the 8-cell stage into single b4.2 blastomeres (Figure 3C). Because the only neural progeny of b4.2 cells are midline neural cells, this results in the selective inhibition of Cad2 expression in midline neural cells (Figure 3C). Inhibiting Cad2 expression in midline neural cells caused a sharp decrease in myosin activity, and a significant increase in boundary roughness, along Ne/Epi boundaries, relative to un-injected sides of the same embryos (Figure 3G and Movie S2, quantified in Figure 3H and 3I). In contrast, myosin activity was significantly increased, and boundary roughness was significantly reduced, along Ne/Ne boundaries between MO-receiving and non- receiving neural cells (Figure 3G and Movie S2; quantified in Figure 3H and 3I), suggesting that creating an ectopic Cad2 expression boundary within neural territory is sufficient to induce myosin activation and increase tension along that boundary. Importantly, zipper progression and neural tube closure were strongly reduced in Cad2 MO-injected embryos (Figure 3J, 3L and 3M). Moreover, the residual zippering was associated with straightening and shortening of the Ne/Ne boundary between normal and Cad2 MO-injected neural cells, suggesting that it is due to active shortening of the ectopic boundary, rather than to a residual contribution of the normal Ne/Epi boundary (Movie S3). We observed similar effects with two different morpholinos (MO1 and MO2), suggesting that these effects are specific to Cad2 inhibition (Figure S2A-S2C). Thus inhibiting Cad2 expression in midline neural cells is also sufficient to block Myosin activation along the Ne/Epi boundary, zipper progression and neural tube closure.

### Differential expression of Cad2 in anterior neural cells directs myosin activation and purse string closure of the anterior neural tube

Following zippering and closure of the posterior nerve tube, the anterior neural tube closes to form the sensory vesicle. Interestingly, we found that anterior neural tube closes through a combination of “zippering” and “purse string” mechanisms, in which activated myosin accumulates along the entire anterior Ne/Epi boundary, and all Ne/Epi junctions shorten simultaneously (Figure 4A, 4B and Movie S4). Interestingly, *Cad2* is expressed in the anterior neural plate before the onset of anterior closure (Figure 4C), suggesting that differential expression of Cad2 across the anterior Ne/Epi boundary controls myosin activation and purse-string closure of the sensory vesicle. To test this, we injected Cad2 MO into the a4.2 blastomere, whose neural progeny are the sensory vesicle precursor cells (Figure 4D). To mark the affected neural cells, we coinjected a marker for neural expression (pZicL⸬Lifeact⸬GFP; Figure S1A and S1B). Injection of Cad2 MO into a4.2 blastomeres sharply decreased myosin activity along the anterior Ne/Epi boundary, relative to the un-injected side of the same embryos (Figure 4E, quantified in Figure 4F), and completely abolished the formation and shortening of a contractile purse string on the injected side. In contrast, myosin activity was significantly increased along the boundary between MO-injected and un-injected anterior neural cells (Figure 4E, quantified in Figure 4F). Thus, differential expression of Cad2 across the Ne/Epi boundary directs myosin activation and boundary contraction during multiple distinct steps in neural tube closure.

**Figure 4.**
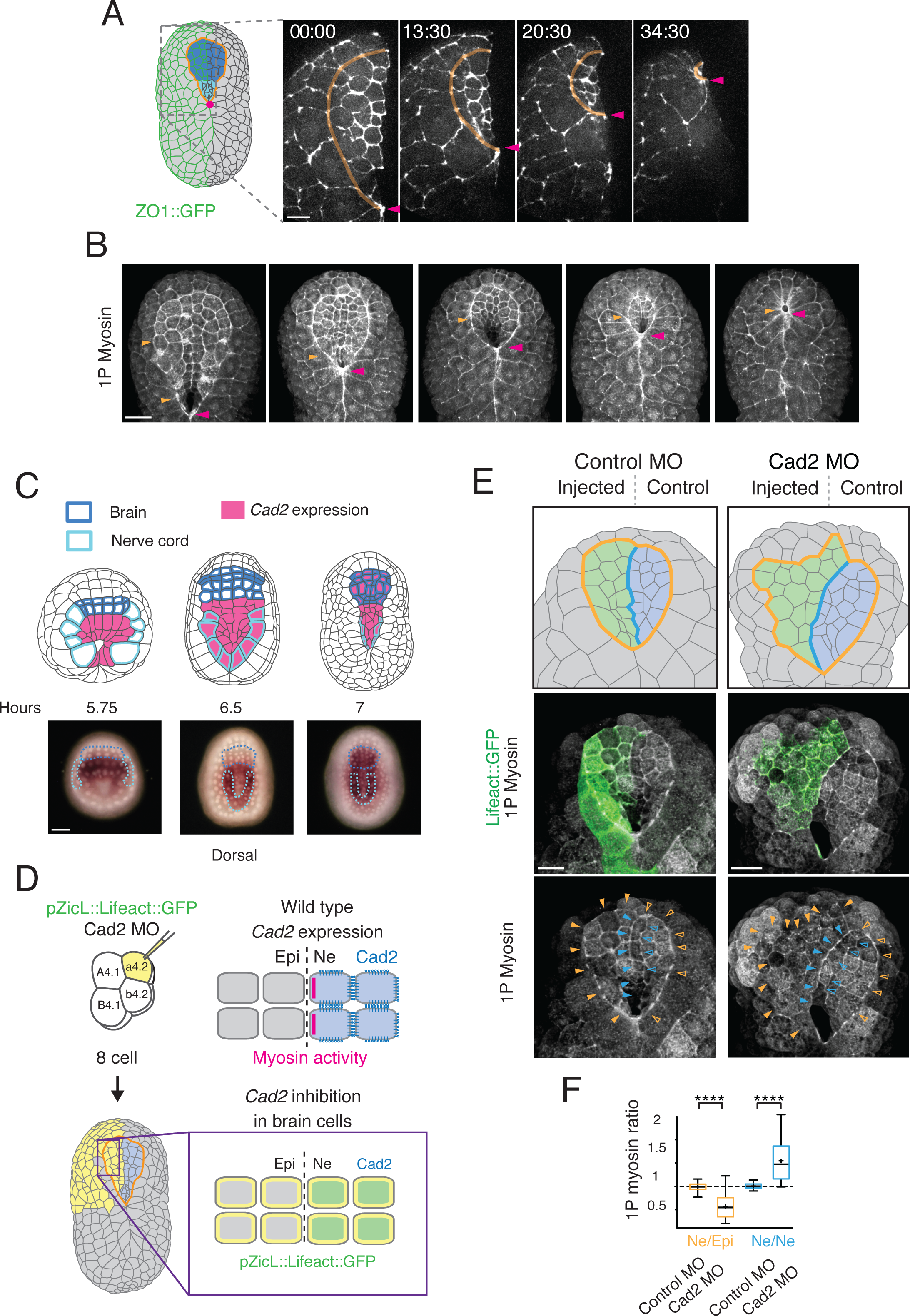
Tissue-specific expression of *Cad2* directs myosin activation and anterior neural tube (brain) closure. (A) Sequence of frames form Movie 4 showing the progression of anterior neural tube (brain) closure in an embryo expressing ZO-1⸬GFP in anterior neural and epidermal cells, driven by the FOG promoter ((Pasini et al., 2006), Figure S1). Magenta arrowheads indicate zipper position. Orange line indicates Ne/Epi boundary. Schematic on left indicates expression domain. (B) Distribution of 1P-myosin at successive steps during brain closure. Orange arrowheads indicate Ne/Epi boundary; Magenta arrowheads indicate zipper position. (C) Cad2 gene expression pattern at late gastrula, early neurula (during nerve cord zippering) and late neurula (during brain closure) stages. Nerve cord cells are outlined in light blue; anterior brain precursors are outlined in dark blue. Scale bars = 25 μm. (D) Co-injection of Cad2 MO with an expression marker (pZicL⸬Lifeact⸬GFP) into a single a4.2 blastomere at the 8-cell stage. Yellow indicates all a4.2 descendants; green indicates anterior neural descendants that received the MO. (E) Effects of Cad2 knockdown in anterior neural cells. Top panel shows schematic dorsal surface view of late neurula stage embryos that were injected with control MO (left) or Cad2 MO (right), then fixed and immunostained for 1P myosin. Yellow indicates neural cells inheriting co-injected morpholino and GFP marker (green) over endogenous Cad2 (blue). Orange line indicates the original Ne/Epi boundary. Blue line indicates boundary between injected and non-injected neural cells. Middle panel shows superposition of GFP marker and 1P-myosin in the same embryos. Bottom panel shows 1P-myosin alone. Open and filled arrowheads indicate corresponding boundaries on injected and non-injected sides of the embryo. Scale bars = 5 μm. (F) Box plots showing 1P myosin ratios for the indicated junction types. Ne/Epi ratios were measured on injected:non-injected junctions. Ne/Ne ratios were measured as the average intensity on junctions between injected and non-injected neural cells (filled blue arrowheads in E) divided by average intensity on junctions between non-injected neural cells (open blue arrowheads in E). Control measurements were made on embryos injected with control morpholino (n > 29 junctions, from > 7 embryos for each condition; **** p < 0.0005, Student’s t test).

### Differential expression and homotypic enrichment of Cad2 directs RhoA activity to the Ne/Epi boundary

We previously reported that the RhoA/Rho kinase signaling is required for myosin activation along the Ne/Epi boundary during neural tube closure (Figure 5A) (Hashimoto et al., 2015). To test whether RhoA activity is spatially patterned in midline neural cells, we constructed the ascidian analogue of a previously validated biosensor for active RhoA by fusing GFP to the RhoA binding domain of Ciona Anillin (henceforth GFP⸬AHPH; (Munjal et al., 2015; Piekny and Glotzer, 2008; Priya et al., 2015; Tse et al., 2012)). To characterize the localization of active RhoA in neural cells during zippering, we expressed GFP⸬AHPH in neural cells using the Etr1 promoter. We observed a local enrichment of GFP⸬AHPH along all Ne/Epi junctions ahead of the zipper, with highest accumulation just ahead of the zipper, as previously observed for myosin (Figure 5B, Movie S5, (Hashimoto et al., 2015)). Significantly, we also observed a rapid decay of the GFP⸬AHPH signal just behind the zipper where heterotypic Ne/Epi contacts are replaced by homotypic Ne/Ne contacts, suggesting that differential expression and homotypic enrichment of Cad2 dynamically controls the local activation of RhoA (Figure 5B and Movie S5).

**Figure 5.**
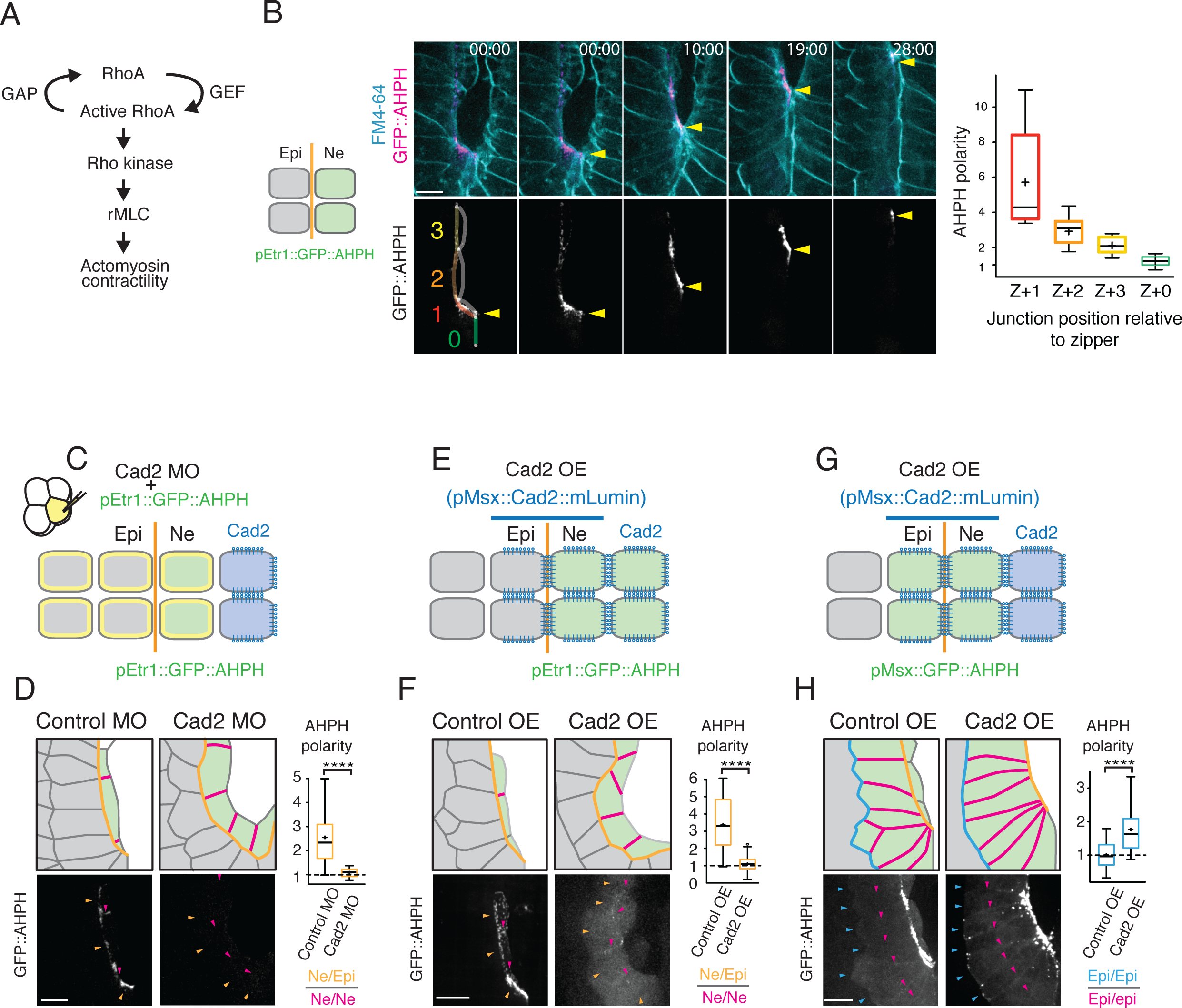
Differential expression of *Cad2* directs polarized activation of RhoA in midline neural cells. (A) RhoA/Rho kinase signaling pathway for myosin activation. (B) A sequence of images (from Movie S5) showing the dynamic distribution of a RhoA biosensor GFP⸬AHPH. Top row shows the biosensor (magenta) and a membrane marker (FM-464; blue); bottom row shows the biosensor alone. Yellow arrowheads indicate zipper position. Right: Quantitation of average GFP⸬AHPH polarity in neural cells as a function of cell position relative to zipper, indicated by color overlays in the first movie frames in left schematic. Box plots show the distribution of average polarity values measured for individual junctions during the period in which an Ne/Epi junction next to the zipper (z + 1) starts and finishes its contraction. (C, D) Effect of Cad2 MO on distribution of GFP⸬AHPH in neural cells. (E, F) Effects of ectopic expression of Cad2 in all midline cells on distribution of GFP⸬AHPH in midline neural cells. (G, H) Effects of ectopic expression of Cad2⸬mLumin in all midline cells on distribution of GFP⸬AHPH in midline epidermal cells. Green: cells expressing GFP⸬AHPH, yellow: cells receiving Cad2 MO, blue: distributions of Cad2 protein. In (D, F), Orange lines and arrowheads indicate midline Ne/Epi junctions. Magenta lines and arrowheads indicate midline Ne/Ne junctions. In (H), Orange lines indicate midline Ne/Epi junctions, blue lines and arrowheads indicate junctions along the ectopic Cad2⸬mLumin expression boundary between midline and non-midline epidermal cells, magenta lines and arrowheads indicate junctions between midline epidermal cells. In (D,F,H) graphs show GFP⸬AHPH polarity measured as ratio of intensities on junction types indicated below each graph. As a control in (D), Control MO was injected into a4.2. As a control in (F,H), Lifeact⸬mLumin was expressed in neural cells. (n > 23 junctions in C, F and H) **** p < 0.0005, by Student’s t test. Scale bars = 5 μm.

To test whether Cad2 controls the spatial pattern of RhoA activity in midline cells, we co-injected Cad2 MO with DNA encoding neural expression of the Rho biosensor (pEtr1⸬GFP⸬AHPH) into single b4.2 cells at the 8-cell stage (Figure 5C). Because we could not directly compare GFP⸬AHPH levels in injected and control sides of the same embryos, we quantified GFP⸬AHPH polarity, defined as the mean GFP⸬AHPH intensity along Ne/Epi junctions divided by half the mean intensity along Ne/Ne junctions (Ne/Epi⸬Ne/Ne intensity ratio; see methods for details). GFP⸬AHPH polarity was completely abolished in midline neural cells descended from b4.2 cells injected with Cad2 MO, relative to embryos injected with a control morpholino (Figure 5D, Movie S6). The loss of GFP⸬AHPH signal along the Ne/Epi boundary in Cad2 MO-injected midline neural cells was not due to absence of GFP⸬AHPH expression, because GFP⸬AHPH was similarly enriched at cleavage furrows in both control and Cad2 MO injected embryos (Figure S3, Movie S6). In complementary experiments, we found that over-expressing Cad2⸬mLumin (variant mKate: (Chu et al., 2009)) in all midline cells (Figure 5E) also abolished GFP⸬AHPH polarity in neural midline cells, relative to embryos expressing a control construct (Figure 5F). Thus, equalizing either high or low levels of Cad2 expression across the Ne/Epi boundary leads to loss of RhoA activity along the Ne/Epi boundary.

Finally, in embryos over-expressing Cad2⸬mLumin in all midline cells (Figure 5G), GFP⸬AHPH was enriched on junctions between Cad2 expressing and non-expressing epidermal cells, relative to the corresponding junctions in control embryos (Figure 5H; enrichment measured as mean Epi/Epi[heterotypic]⸬Epi/Epi[homotypic] intensity ratio; see methods for details). Thus differential expression and homotypic enrichment of Cad2 is sufficient to promote local activation of RhoA along ectopic expression boundaries.

To summarize thus far, we find that abolishing homotypic enrichment of Cad2 in midline neural cells, through either uniform enrichment or uniform depletion, inhibits RhoA and myosin activity along the Ne/Epi boundary. On the other hand, inducing ectopic homotypic enrichment of Cad2 induces activation of RhoA and myosin. Thus it is not the presence or absence of Cad2 *per se* that controls RhoA and myosin activity. Rather, it is the enrichment of Cad2 on homotypic contacts that directs RhoA and myosin activity to heterotypic contacts within the same cells.

### Cad2 directs polarized enrichment of Gap21/23 in midline neural cells

How does homotypic enrichment of Cad2 direct activation of RhoA and Myosin II to Ne/Epi boundaries? One possibility is that Cad2 sequesters an inhibitor of RhoA to Ne/Ne junctions, preventing it from accumulating on Ne/Epi junctions. To explore this possibility, we focused on Rho GTPase activating proteins (RhoGAPs), which inhibit Rho GTPase signaling by inducing GTP hydrolysis (Bos et al., 2007). We screened the Ciona genome for putative RhoGAPs, whose orthologues localize to cell-cell junctions in other organisms, and which are homotypically enriched on Ne/Ne junctions when expressed in neural cells using the pEtr1 promoter. One of the five RhoGAPs we tested (Gap5, Gap11A, Gap21/23, Gap22/24/25, Gap29; see Figure S4A and S4B), only Gap21/23 was homotypically enriched in neural cells. Gap21/23 is the Ciona orthologue of Human RhoGAPs ARHGAP-21 and ARHGAP-23 (Peck et al., 2002) (previously known as ARHGAP-10 (Bassères et al., 2002)) (Figure S4A). Using in situ hybridization, we determined that *Gap21/23* mRNA is expressed in all embryonic cells until the 32-cell stage; it decays before the onset of gastrulation, and then it is expressed again, specifically in posterior midline epidermal and neural cells, just before the onset of zippering (Figure 6A and S4C). When expressed using the Etr1 promoter, GFP⸬Gap21/23 was enriched on Ne/Ne junctions, including newly-formed Ne/Ne junctions behind the zipper, and virtually absent from Ne/Epi junctions during zippering (Figure 6B, S6B), suggesting that Cad2 may pattern RhoA/myosin activity by localizing Gap21/23.

**Figure 6.**
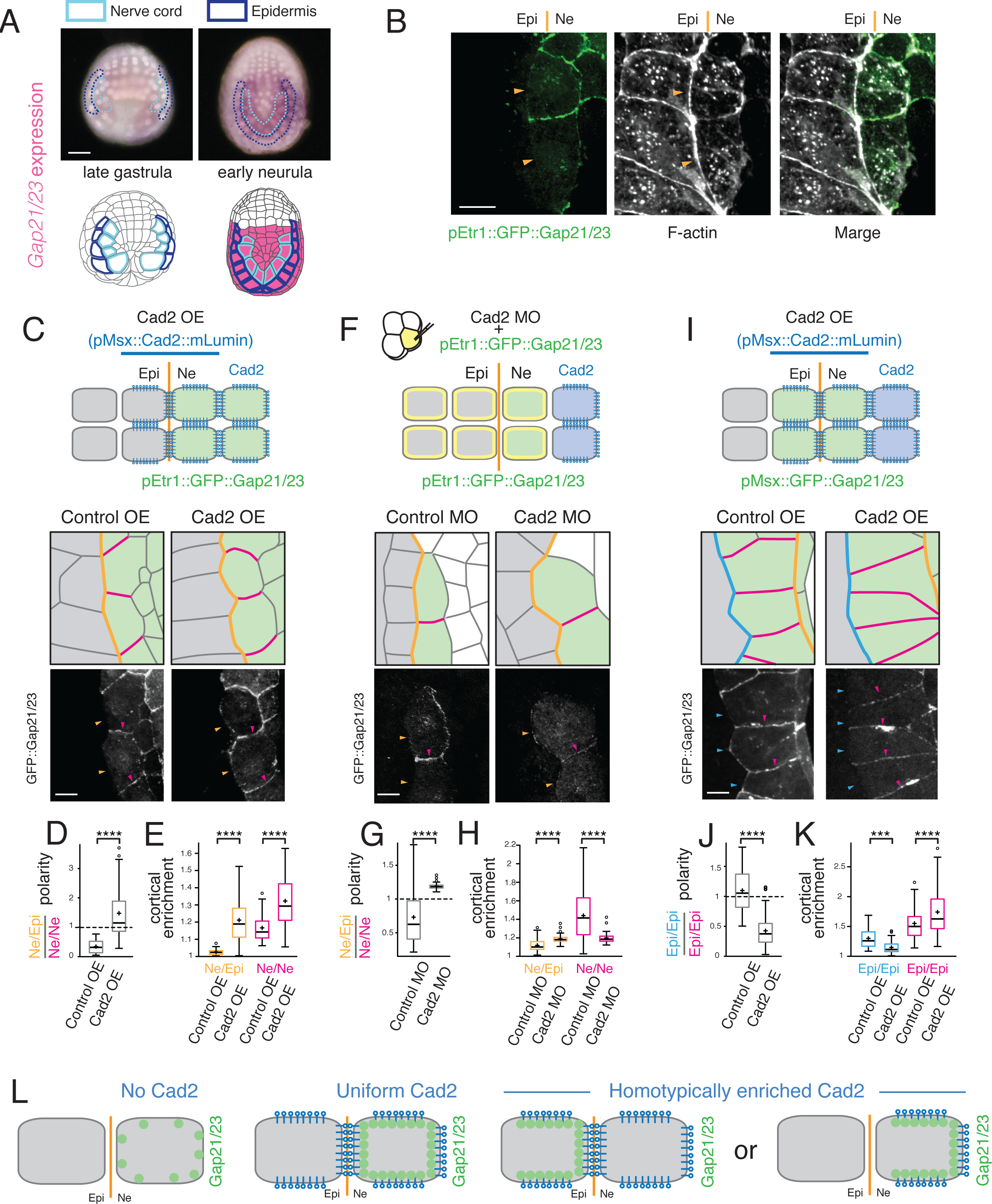
Sequestration by homotypically enriched Cad2 polarizes Gap21/23. (A) *Gap21/23* expression (magenta) at late gastrula and early neurula stages. Light blue and dark blue indicate midline neural and epidermal cells respectively. Scale bars = 25 μm. (B) Localization of GFP⸬Gap21/23 in neural cells. Embryo was fixed and co-stained with phalloidin. Orange lines and arrowheads indicate Ne/Epi boundary. Scale bars = 5 μm. (C-K) Distributions of GFP⸬Gap21/23 for different expression patterns of Cad2. Scale bars, 5 μm. (C-E) Effects of ectopically expressing Cad2⸬mLumin in all midline cells on distribution of GFP⸬Gap21/23 in midline neural cells. (F-H) Effects of knocking down Cad2 on distribution of GFP⸬Gap21/23 in midline neural cells. (I-K) Effects of ectopically expressing Cad2⸬mLumin in all midline cells on distribution of GFP⸬Gap21/23 in midline epidermal cells. (C, F) Orange lines and arrows indicate Ne/Epi junctions, and magenta lines and arrows indicate Ne/Ne junctions, in midline neural cells. (I) Magenta lines and arrowheads indicate junctions between midline epidermal cells. Blue lines and arrowheads indicate junctions between midline and non-midline epidermal cells. (D, G, J) GFP⸬Gap21/23 polarity measured as ratio of intensities on junction types indicated below each graph. (E, H, K) relative enrichment of GFP⸬Gap21/23, measured as the ratio of junctional to cytoplasmic intensity for junction types indicated below each graph. As a control in C-E, I-K, Lifeact⸬GFP was expressed in neural cells. As a control in F-H, control MO was injected into a4.2. (n > 23 junctions, from > 9 embryos for each of the conditions in D, E, G, H, J and K) *** p < 0.005, **** p < 0.0005,, Student’s t test. (L) Schematic summary of Gap21/23 distributions for different Cad2 expression and localization patterns.

To test whether Cad2 controls Gap21/23 localization, we manipulated Cad2 expression patterns and measured changes in GFP⸬Gap21/23 polarity (measured as the ratio of intensity on Ne/Epi vs Ne/Ne boundaries) and junctional enrichment (measured as the ratio of junctional⸬cytoplasmic intensities) in midline neural cells (see Methods for details). Over-expressing Cad2 in all midline cells abolished GFP⸬Gap21/23 polarity in midline neural cells (Figure 6C and 6D); this loss of polarity was accompanied by increased enrichment of GFP⸬Gap21/23 on both Ne/Epi and Ne/Ne junctions (Figure 6E), consistent with increased recruitment of Gap21/23 by uniformly enriched Cad2. Inhibiting Cad2 expression in midline neural cells also abolished GFP⸬Gap21/23 polarity (Figure 6F and 6G), but in this case, the loss of polarity was associated with decreased enrichment on Ne/Ne junctions and increased enrichment on Ne/Epi junctions (Figure 6H), suggesting that Cad2-independent mechanisms can recruit Gap21/23 to equal levels on all junctions, and that homotypically-enriched Cad2 can bias recruitment away from Ne/Epi junctions and towards Ne/Ne junctions.

To test this further, we used the midline promoter pMsx to drive ectopic expression of GFP⸬Gap21/23 in midline epidermal cells, which do not normally express Cad2 (Figure 6I). When ectopically co-expressed with a control protein (Lifeact⸬GFP) in midline epidermal cells, GFP⸬Gap21/23 was uniformly enriched on all Epi/Epi junctions (Figure 6I and 6K). In contrast, ectopic co-expression with Cad2 induced polarized enrichment of GFP⸬Gap21/23 in midline epidermal cells (Figure 6I and 6J), and again, this gain of polarity was associated with increased enrichment of GFP⸬Gap21/23 on homotypic (Cad2/Cad2) junctions, and decreased enrichment on heterotypic (Cad2/-) junctions (Figure 6K). Thus, in the absence of Cad2, Gap21/23 can accumulate at uniformly low levels on all junctions (Figure 6L, left), uniformly enriched Cad2 induces uniformly high levels of GAP-21/23 accumulation (Figure 6L, middle), and homotypically enriched Cad2 induces polarized enrichment of Gap21/23 by recruiting Gap21/23 to homotypic junctions and away from heterotypic junctions (Figure 6L, right).

### Gap21/23 is required for polarized activation of RhoA and myosin II, zippering, and neural tube closure

To assess functional contributions of Gap21/23 to polarized activation of myosin II, zippering and neural tube closure, we first examined the consequences of overexpressing Gap21/23 in midline neural cells. Embryos over-expressing GFP⸬Gap21/23 in all neural cells (using the Etr1 promoter) developed normally up to the zippering stage, but then failed to undergo normal zippering (Figure S5A, S5B). Over-expressed GFP⸬Gap21/23 was homotypically enriched in midline neural cells, but with detectable accumulation on Ne/Epi junctions (Figure S5F). Importantly, polarized distributions of RhoA (assessed by co-overxpression pEtr1⸬GFP⸬AHPH) and active myosin (assessed by immunostaining 1P myosin) were completely abolished in midline cells over-expressing Gap21/23 (Figure S5C-S5G). Although we could not assess absolute changes in GFP⸬AHPH accumulation, loss of myosin polarity was associated with decreased 1P myosin on both Ne/Epi and Ne/Ne junctions (Figure S5H), relative to the unaffected sides of the same embryos. Thus over-expressing Gap21/23 leads to local accumulation of Gap21/23, and a local decrease in myosin activity, and decrease tension on Ne/Epi junctions (Figure S5I), suggesting that Gap21/23 can act locally at junctions to inhibit RhoA and myosin activity.

Finally, we asked whether inhibiting the expression of Gap21/23 affects polarized activation of RhoA and myosin II, zippering and neural tube closure. To bypass a possible requirement for Gap21/23 during early development, we injected Gap21/23 MO into b4.2 cells at the 8-cell stage (Figure 7A). These embryos developed normally until just before the onset of zippering but then failed to undergo zippering and neural tube closure (Figure 7A and 7B). Polarized enrichment of GFP:AHPH and 1P myosin on Ne/Epi junctions was completely lost in midline neural cells descended from b4.2 cells injected with Gap21/23 MO (confirmed with a second MO; Figure S5I), but not from cells injected with a control MO. (Figure 7C-F, Movie S7). This was accompanied by increased roughness (indicating reduced tension) along Ne/Epi boundary (Figure 7G). Although we could not measure changes in RhoA activity relative to internal controls, the loss of polarized myosin activity was accompanied by decreased 1P myosin on Ne/Epi junctions, and increased 1P myosin on Ne/Ne junctions, relative to the un-injected sides of the same embryos (Figure 7H). Thus, in the absence of Gap21/23, myosin (and likely RhoA) is constitutively and uniformly active on all junctions, and our data suggest that one or more factors required for activation are limiting, such that enrichment of Gap21/23 at Ne/Ne junctions by Cad2 displaces this activity away from Ne/Ne junctions and towards Ne/Epi junctions.

**Figure 7.**
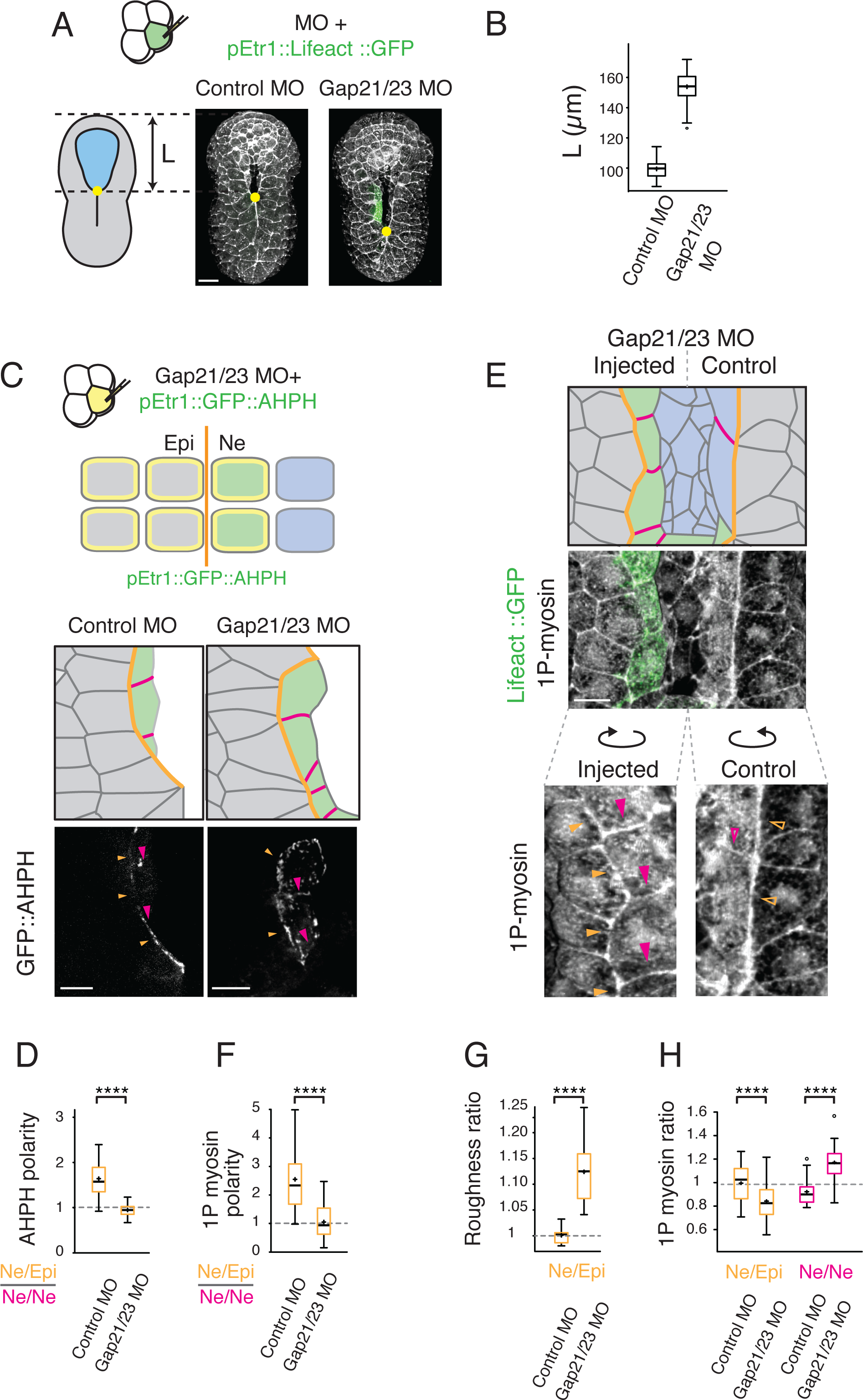
Gap21/23 is required for polarized activation of RhoA/myosin and zipper progression. (A) Inhibition of zipper progression in Gap21/23 knockdown embryos. Left schematic shows measurement of “un-zippered length” L. Microinjected embryos expressing neural-specific co-injection marker (pEtr1⸬Lifeact⸬GFP) and fixed and co-stained with phalloidin. Scale bars, 10 μm. (B) Quantification of zipper progression in Gap21/23 knockdown embryos (n > 17 embryos for each condition). (C, D) Effects of Gap21/23 knockdown on distribution of GFP⸬AHPH in midline neural cells. Top schematic: yellow indicates cells receiving Gap21/23 MO; green indicates cells expressing pEtr1⸬GFP⸬AHPH. Bottom panels: Orange lines and arrowheads indicate Ne/Epi junctions. Magenta lines and arrowheads indicate Ne/Ne junctions in midline neural cells. Scale bars = 5 μm. (D) GFP⸬AHPH polarity measured as the ratio of intensities on Ne/Epi⸬Ne/Ne junctions. (E, F, H) Effects of Gap21/23 knockdown on distribution of 1P-myosin in midline neural cells. (E) Top panel shows dorsal surface view of a neurula stage embryo, injected as shown in (A), then fixed and immunostained for 1P myosin. Color fill indicates neural cells inheriting co-injected morpholino and GFP marker (green) over endogenous Cad2 (blue). Orange lines and arrowheads indicates the original Ne/Epi boundary. Magenta lines and arrowheads indicate Ne/Ne junctions in midline neural cells. Second panel: superposition of GFP marker and 1P-myosin in the same embryo. Third panel: 1P-myosin alone. Open and filled arrowheads indicate corresponding boundaries on injected and non-injected sides of the embryo. Scale bars = 5 μm. (G) Roughness ratio in Gap21/23 knockdown embryos. (n > 9 embryos for each condition) (H) 1P myosin ratio in Gap21/23 knockdown embryos (n > 14 junctions from > 9 embryos for each condition in D, F, and H). **** p < 0.0005, Student’s t test.

## Discussion

Embryos use dynamic control of actomyosin contractility to maintain tissue boundaries and drive tissue morphogenesis (reviewed in (Dahmann et al., 2011; Fagotto, 2015; Heisenberg and Bellaiche, 2013; Lecuit et al., 2011)), but how they implement this control at the cellular level, and how it is coordinated with local tissue remodeling, remain poorly understood. Here we describe how tissue-specific expression of Cad2 in neural cells patterns RhoA/myosin II activity to create a dynamic tissue-level contractile asymmetry that is essential for zippering and neural tube closure. Our data support a model in which: (a) Differential expression of Cad2 drives its homotypic enrichment in midline neural cells, (b) homotypically enriched Cad2 sequesters Gap21/23 to neural cell contacts and away from the Ne/Epi boundary, and (c) polarized Gap21/23 redirects RhoA/Myosin II activity away from homotypic junctions and towards heterotypic junctions. Local activation of RhoA/Myosin II at heterotypic Ne/Epi junctions ahead of the zipper, and rapid inactivation at newly-formed homotypic Ne/Ne junctions behind the zipper, supports a self-adjusting tissue-level contractile asymmetry that is required for unidirectional zipper progression and neural tube closure.

### Differential expression directs homotypic enrichment of Cad2 in midline neural cells

In wild type embryos, Cad2 is expressed in the neural primordium, it is highly enriched on homotyopic Ne/Ne contacts, and absent from heterotypic Ne/Epi contacts. In contrast, when Cad2 is over-expressed in all midline cells, it is enriched at all homotypic contacts between Cad2 expressing midline cells, regardless of cell identity (Ne/Ne, Ne/Epi, or Epi/Epi), and absent from heterotypic Epi/Epi contacts. Thus differential expression of Cad2 dictates its homotypic enrichment, independent of the underlying cell identities. Previous cell-mixing experiments have documented a tendency for cadherins (including VE-Cadherin, the human orthologue of Cad2) to become enriched at boundaries between expressing cells, (but not between expressing and non-expressing cells) (Hirano et al., 1987; Klompstra et al., 2015; Navarro et al., 1998). Here we show that the same rules operate for Cad2 within an intact embryonic epithelium. The underlying mechanism(s) remain unclear, but likely involve multiple forms of feedback downstream of trans-homophillic engagement, including local stabilization of clustered Cadherins at cell-cell contacts (Cavey et al., 2008; Foote et al., 2013; Nose et al., 1988), and local inhibition of endocytosis (Izumi et al., 2004).

### Gap21/23 is the key intermediary between homotypically enriched Cad2 and polarized activation of RhoA/Myosin II

Three observations identify Gap21/23 as the key intermediary between Cad2 and polarized activation of RhoA/Myosin II. First, Gap21/23 is expressed in neural cells from just before the onset of zippering, and inhibiting its expression in neural cells blocks zippering and neural tube closure. Second, homotypically enriched Cad2 dictates polarized enrichment of Gap21/23 in midline neural cells. When GFP⸬Gap21/23 is expressed in the neural primordium in otherwise wildtype embryos, it is co-enriched with Cad2 in midline neural cells at Ne/Ne, but not at Ne/Epi junctions. Inhibiting Cad2 expression in neural cells leads to uniformly weak accumulation of GFP⸬Gap21/23, with *decreased* enrichment on Ne/Ne junctions and *enhanced* enrichment at Ne/Epi junctions. In contrast, forcing uniformly high enrichment of Cad2 in midline neural cells leads to uniformly strong accumulation of GFP⸬Gap21/23 on all junctions, with enhanced enrichment on *both* Ne/Epi and Ne/Ne junctions. These data are consistent with a model in which a limiting pool of Gap21/23 can associate weakly with all cell contacts independently of Cad2, and homotypically enriched Cad2 sequesters Gap21/23 to Ne/Ne junctions, preventing its association with Ne/Epi junctions.

The molecular basis for sequestration of Gap21/23 by Cad2 remains unclear. E-cadherin acts through a-catenin and p120-catenin to recruit Gap21/23 orthologues to cell-cell contacts in early *C. elegans* embryos (Klompstra et al., 2015) and in vertebrate tissue culture (Sousa et al., 2005). The Cad2 orthologue VE-cadherin acts through p120-catenin to recruit other GAPs, such as p190 RhoGAP (Zebda et al., 2013). Thus an interesting possibility is that Cad2 uses a-catenin and p120-catenin to recruit Gap21/23 to Ne/Ne contacts in ascidian embryos, although this remains to be tested.

Finally, Gap21/23 is required to polarize RhoA and Myosin II activity in midline neural cells. In Gap21/23-depleted neural cells, RhoA/Myosin II activity is uniformly high on all junctions, suggesting that upstream activators are uniformly enriched in neural cells and that local inhibition of RhoA by Gap21/23 dictates its polarized activity. *In vitro*, ARHGAP21 can act as a GAP towards either RhoA or CDC-42, with preference for the latter (Dubois et al., 2005) Likewise, in culture or *in vivo*, ARHGAP21 and its orthologues inhibit either RhoA or CDC-42, depending on cell type and subcellular localization (Barcellos et al., 2013; Klompstra et al., 2015; Lazarini et al., 2013; Marston et al., 2016; Zhang et al., 2016). Thus, we currently favor the simplest possibility that Gap21/23 inhibits RhoA directly by stimulating GTP hydrolysis.

Although we have focused on the control of RhoA/myosin activity in midline neural cells, active RhoA and myosin are also enriched along the Ne/Epi boundary in midline epidermal cells (Figure 1C, D, and data not shown). Thus additional, and possibly analogous, mechanism(s) may operate in epidermal cells. Alternatively, epidermal cells could activate myosin II in response to tension produced by local contraction of neighboring neural cells (Fernandez-Gonzalez et al., 2009).

### Comparison to other systems: Parallels and differences

Tissue-specific Cadherin expression has long been thought to play a major role in the self-organization of embryonic tissues (Takeichi, 1995), but the underlying mechanisms have remained poorly understood (Fagotto, 2015). Our study provides the first clear example in chordates of a mechanism by which tissue-specific expression of a classic cadherin localizes actomyosin contractility to a tissue boundary to control morphogenesis. However, recent work in other contexts points to a more general role for differential expression of cadherins, and other homphilic binding proteins, in localized control of myosin II at heterotypic cell contacts. In early *C. elegans* embryos, E-cadherin recruits the orthologue of Gap21/23 (PAC-1) to all somatic cell contacts, where it inhibits CDC-42, restricting CDC-42 activity to non-contacting (apical) cell surfaces (Anderson et al., 2008; Klompstra et al., 2015). During gastrulation, endoderm-specific signal(s) couple apical CDC-42 activity to the local recruitment of a CDC-42 effector MRCK-1 to activate myosin II and drive endoderm cell ingression (Marston et al., 2016). In the developing Drosophila eye, differential expression and homotypic enrichment of N-cadherin directs myosin activity to heterotypic junctions between cone and primary pigment cells. In this system, N-Cadherin may not act by sequestering an inhibitor of Myosin II to homotypic contacts. Instead, a small amount of unbound N-Cadherin appears to promote contractility at heterotypic contacts, although the mechanism remains unknown (Chan et al., 2017).

Looking further afield, it has been proposed that Drosophila Echinoid localizes myosin II to the epidermis/amnioserosa boundary during dorsal closure by sequestering PAR-3/Bazooka (Laplante and Nilson, 2011), although direct evidence for this mechanism is still lacking. During salivary gland invagination, the intracellular domain of Crumbs recruits atypical protein kinase C (aPKC) to homotypic contacts between Crumbs expressing cells, where it is thought to phosphorylate and inhibit Rho Kinase (Ishiuchi and Takeichi, 2012), restricting accumulation of Rho kinase and myosin activity to the boundary of the salivary gland rudiment (Röper, 2012). In summary, there seem to be many ways in which embryos couple different homophilic binders to different intermediate signaling pathways to localize myosin activity to heterotypic tissue boundaries, and it will be important to detail the essential intermediate steps for individual examples, as we have done here, before it will be possible to draw more general conclusions.

### Implications for dynamic control of zipper progression in ascidians, and other chordates

We previously showed that zipper progression involves two modes of control over myosin II (Hashimoto et al., 2015): Sequential activation of myosin II and rapid contraction of individual Ne/Epi junctions just ahead of the zipper provides the power stroke for zipper progression; higher activity of myosin II on all Ne/Epi junctions ahead of the zipper, and lower activity on newly formed Ne/Ne and Epi/Epi junctions just behind the zipper, creates a tissue-level imbalance of force that converts these sequential contractions efficiently into net forward movement of the zipper (Hashimoto et al., 2015).

Our current findings now explain how this tissue level force imbalance is dynamically maintained as the zipper progresses (Figure 8): Ahead of the zipper, sequestration of Gap21/23 and polarization of RhoA/Myosin II leads to local higher levels of myosin activity and tension along the Ne/Epi boundary. Behind the zipper, rapid engagement of Cad2 at newly formed Ne/Ne contacts leads to rapid recruitment of Gap21/23, local inhibition of RhoA/myosin II, and reduced tension along the Ne/Ne boundary. These observations imply a tissue-level feedback loop in which local remodeling of tissue contacts is rapidly converted to local changes in force production to maintain a tissue-level force imbalance (higher ahead of the zipper; lower behind) that converts rapid all-or-none contractions ahead of the zipper into efficient net forward movement.

**Figure 8.**
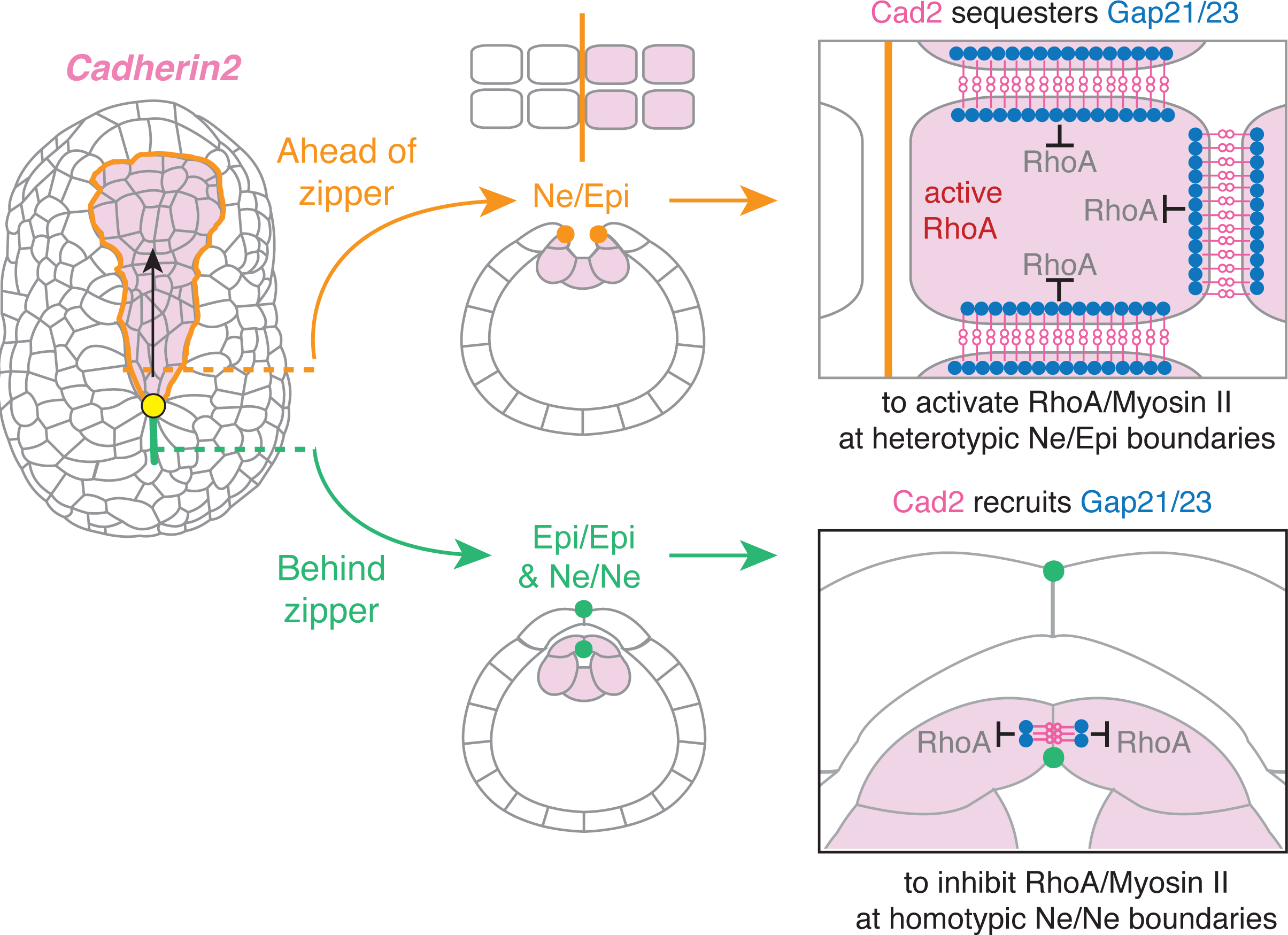
Model for self-adjusting tissue level control of contractile asymmetry. Ahead of the zipper, homotypically enriched Cad2 sequesters Gap21/23 to Ne/Ne junctions, directing RhoA/myosin II activity to Ne/Epi junctions. Behind the zipper, Cad2 recruits Gap21/23 to newly formed Ne/Ne junctions, inhibiting RhoA and myosin II at those junctions (see text for details).

Interestingly, we have found that in addition to abolishing tissue level contractile asymmetries, equalizing Cad2 expression across the Ne/Epi boundary (or inhibiting Gap21/23 in midline neural cells) also blocks strong activation of RhoA and myosin II just ahead of the zipper and completely blocks zipper progression. Thus local signaling by homotypically-enriched Cad2 is also required for the strong sequential activation of RhoA/myosin II just ahead of the zipper that provides the power stroke for zipper progression. The nature of this requirement, and how strong contractions are localized to a zone just ahead of the zipper, remains to be determined.

Recent studies of zippering and neural tube closure in mouse embryos suggest that an actomyosin cable forms between epithelial and neuroectodermal cells, and contributes to forces that drive zippering (Galea et al., 2017). Thus it will be interesting to determine whether mechanisms similar to those we describe here also pattern myosin activity during neural tube closure in mouse embryos.

## Material and Methods

### Embryo culture

*Ciona robusta* adults were collected and shipped from Half Moon Bay, Oyster Point and San Diego (M-Rep, CA) and then maintained in oxygenated sea water at ~ 16°C. Fertilization, staging, dechorionation and electroporation were conducted as previously described (Bertrand et al., 2003; Corbo et al., 1997; Hotta et al., 2007). We cultured embryos in 5-cm plastic petri dishes coated with 1% agarose and filled with HEPES-buffered artificial seawater (ASWH) (Pasini et al., 2006).

### Constructs for tissue-specific expression of FP fusions

To create Gateway destination vectors for tissue-specific expression of FP fusions, we first amplified *GFP* and *mLumin* (Chu et al., 2009), digested with *Stu1*, *or EcoRV and BeglII* together, and then inserted FP-encoding sequences into the 5’ or 3′ entry site of a standard Gateway pFOG⸬RfA cassette (Roure et al., 2007); FOG (Friend of GATA) is also known as Zfpm (Figure S1A), but we refer to it here as FOG, to create Gateway RfA cassettes: pFOG⸬RfA⸬GFP, pFOG⸬GFP⸬RfA pFOG⸬mLumin⸬RfA, and pFOG⸬RfA⸬mLumin. We amplified previously characterized promoter regions of *Epi1*, *Etr1*, *Msx* and *ZicL* from Ciona genomic DNA (Roure et al., 2014; Sasakura et al., 2009; Shi and Mike Levine, 2008; Veeman et al., 2010). We fused the 218 base pair basal promoter of pFOG (Roure et al., 2007) to the C-terminus of the Epi1, Etr1, Msx and ZicL promoters by PCR. We digested the resulting fusions with *Xho1 and Stu1*, then used infusion (Takara) or HiFi DNA assembly (New England BioLab) kits to replace the full FOG promoter (Pasini et al., 2006) in pFOG⸬RfA⸬3xGFP (Hashimoto et al., 2015), pFOG⸬RfA⸬GFP, pFOG⸬GFP⸬RfA pFOG⸬mLumin⸬RfA, or pFOG⸬RfA⸬mLumin.

We amplified full open reading frames of *Cad2* and *Gap21/23* from the unigene collection (Cogenics) clones VES99_P13 and VES100_J09 respectively. We amplified a fragment corresponding to the conserved AHPH domain of *Ciona robusta* Anillin from Cogenics clone VES88_P24. We fused the sequence encoding *Saccharomyces cerevisiae* Lifeact: (Riedl et al., 2008) to the N terminus of GFP. We created Gateway entry clones containing *Cad2*, *Gap21/23*, *Lifeact⸬GFP* and *AHPH*, respectively, using pCR8/GW/TOPO TA Cloning Kits (Invitrogen). We then used the Gateway LR reaction (Invitrogen) to recombine these Gateway entry clones, or an entry clone for iMyo⸬GFP (Hashimoto et al., 2015), into the tissue-specific destination vectors described above. Table S1 lists the primers that we used to amplify specific insert sequences from template clones or Ciona genomic DNA.

**Table S1.**
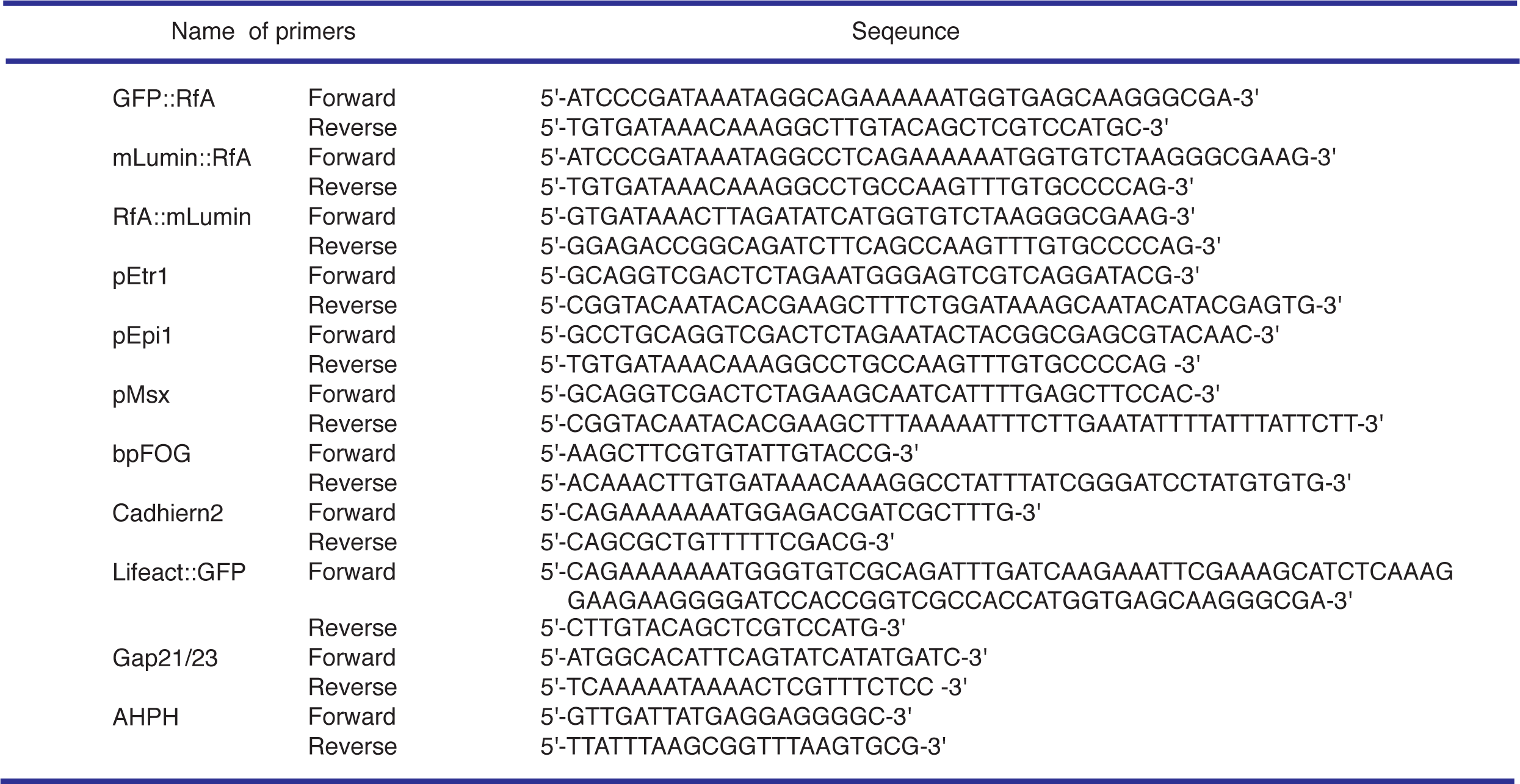
Oligonucleotides Sequence.

### Whole-mount in situ hybridization

We performed detection of *Cad2*, *Gap21/23* mRNAs by whole-mount in situ hybridization with digoxigenin (DIG)-labeled antisense RNA probes as previously described (Wada et al., 1995), using a DIG-RNA labelling kit (Sigma, cat# 11175025910) to synthesis the probes.

### Microinjection of plasmids and MOs

We microinjected eggs as described previously (Bertrand et al., 2003), using 10-25 ng/μl for plasmid DNA, and 0.25–0.5 mM for morpholino oligonucleotides (MO). The sequences of translation-blocking MOs (Gene Tools, LLC) were as follows:

Cad2 MO1, 5′-ATCGTCTCCATGTTGTACTTTATGT-3′

Cad2 MO2, 5′-TTCGCAGTTTTAGTTTCACACTCTT-3′

Gap21/23 MO1, 5′-TGTGCCATTTCAAACACGTTGAAGC-3′

Gap21/23 MO2, 5′-ATTATCTCTTCACAAATCCATTCTT-3′

MO1s cover the starting methionine, and MO2s cover 5′UTR and do not overlap MO1s.

### Immunostaining and imaging of phosphomyosin

We fixed and stained embryos with alexa 488 or 514 phalloidin (ThermoFisher) and a mouse polyclonal antibody raised against ser19-phosphorylated myosin regulatory light chain (1P myosin, Cell Signaling) as previously described (Hashimoto et al., 2015). We collected Z-stacks of confocal images on a Zeiss LSM 880 confocal microscope with a 40×/1.3 oil- immersion objective at 0.5 μm intervals. We rendered 3D projections in ImageJ 3D Viewer (http://rsbweb.nih.gov/ij/plugins/3d-viewer/).

### 4D live time-lapse imaging

We performed 4D live time-lapse imaging as described previously (Hashimoto et al., 2015). Briefly, we settled embryos onto gelatin-formaldehyde coated bottom plates (TED PELLA) filled with seawater at ~18°C. We acquired images using a Nikon ECLIPSE-Ti inverted microscope equipped with 20x and 60x water-immersion lenses, solid-state 50 mW 481 and 561 laser excitation (Coherent), a Yokogawa CSU-X1 spinning disk scan head, an Andor iXon3 897 EMCCD camera, yielding an effective power of 9.1 mW (resp. 14.1 mW). We routinely used 20% of the laser power for imaging on both channels. We collected whole embryo views (resp. magnified views of the zipper region) using 20×/0.75NA (resp. 60×/1.2NA) water-immersion objectives. For some experiments, we labeled cell membranes using the fluorescent lipophilic dye FM4-64 (5μg/ml, Molecular Probes, #T13320). We acquired Z stacks using an x-y motorized stage (for multiple-location imaging) and a fast piezoelectric Z-axis stepper motor with focus steps taken at 0.5 to 1 μm intervals.

### Quantitative analysis of 1P Myosin and fluorescence protein intensities

We performed all image analysis and processing in ImageJ (http://rsbweb.nih.gov/ij/). To quantify junctional intensities of 1P myosin or FP proteins, we constructed maximum intensity projections of image stacks containing the apical surfaces of all relevant cells. We measured the average pixel intensity in three-pixel-wide lines drawn along a junction of interest, excluding regions very close to vertices. Then we subtracted the mean cytoplasmic intensity, measured in junction-adjacent regions, to estimate the junction-specific signal, which we refer to as the average junctional intensity. For mosaically-expressed FP-tagged proteins, we scaled the average junctional intensity between two expressing cells by a factor 0.5 to account for the double contribution. We quantified junctional enrichment as the ratio of average junctional intensity to the mean intensity of junction-adjacent cytoplasmic regions. For 1P myosin ratio, we quantified changes in junctional intensities induced by experimental perturbations as the ratio I_exp_/I_cont_, where I_exp_ the average intensity along a junction on the perturbed side and I_cont_ is the average intensity of the corresponding junction on the contralateral (control) side of the same embryo. We quantified polarity of 1P myosin or FP-tagged proteins as I_AP_/I_ML_, where I_AP_ and I_ML_ are the average intensities for junctions oriented along the anterior-posterior and mediolateral axes, respectively.

### Measurements of iMyo⸬GFP polarity during primary junction contraction

We measured junctional myosin intensity vs time during primary junction contractions in time-lapse sequences acquired at 60s intervals from embryos expressing iMyo⸬GFP. Each frame was the maximum intensity projection of 11 images collected at 0.75 mm steps in z near the apical surface. We only analyzed movies in which zippering was less than 25% complete at the beginning and proceeded to completion. We measured junction lengths manually using ImageJ. To identify the beginning of each primary contraction, we smoothed the junction length data using a 5-frame moving average, and then we determined the time point at which shortening speed increased above a threshold level. We took the end of the contraction to be the time point at which the junction length decreased below a minimal value. We measured junctional intensities as described above for each frame during a primary contraction. Then we aligned data for multiple contractions with respect to the onset of contraction (Fig S1C, D). To measure myosin polarity relative to cell position for a single primary contraction event, we measured the iMyo⸬GFP polarity as described above for cells one, two and three away from the advancing zipper, then we averaged these measurements over the duration of the primary contraction. Figure 1E,F report the mean and standard error for these measurements over many individual primary contraction events.

## Supplemental Figure legends

**Figure S1 (related to Figure 1).**
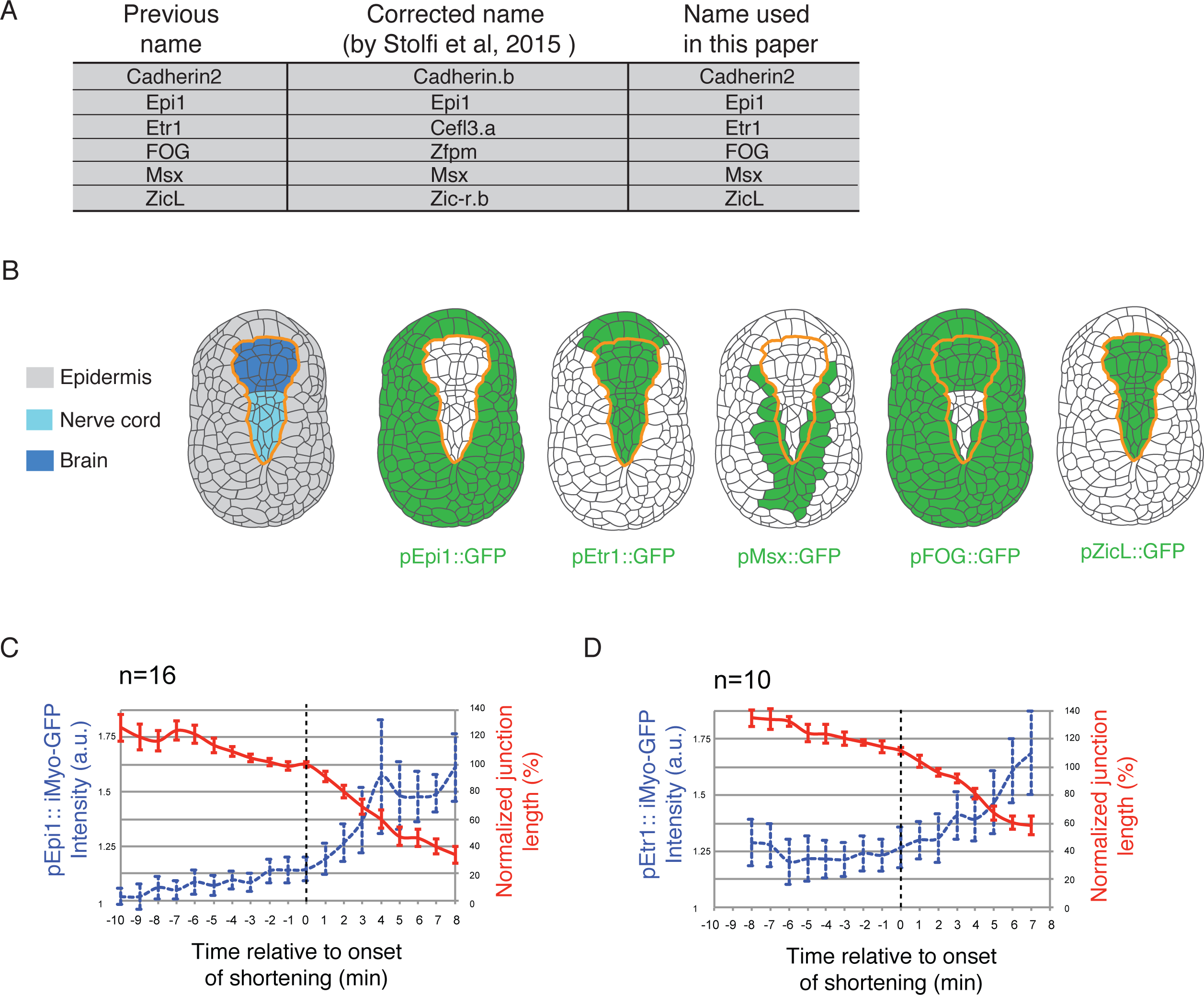
iMyo⸬GFP accumulates during junction shortening events. (A) Name of genes in this study (Stolfi et al., 2015). (B) GFP expression pattern driven by different promoters used in this study. Green cells indicate GFP-expressing cells. Orange line indicates Ne/Epi boundary. (C, D) Relationship between iMyo-GFP intensity and junctional length during individual junction shortening events in embryos iMyo-GFP in epidermal cells (C) or neural cells (D). Data from individual junctions were aligned with respect to the onset of shortening (see Materials and Methods for details). Red lines: normalized average junction length. Blue dashed lines: relative iMyo-GFP fluorescence intensity averaged along the junction, excluding the vertices. n = 16 junctional shortening events from 10 embryos in (C), n = 10 junctional shortening events from 8 embryos in (D). Error bars are SEM.

**Figure S2 (related to Figure 3).**
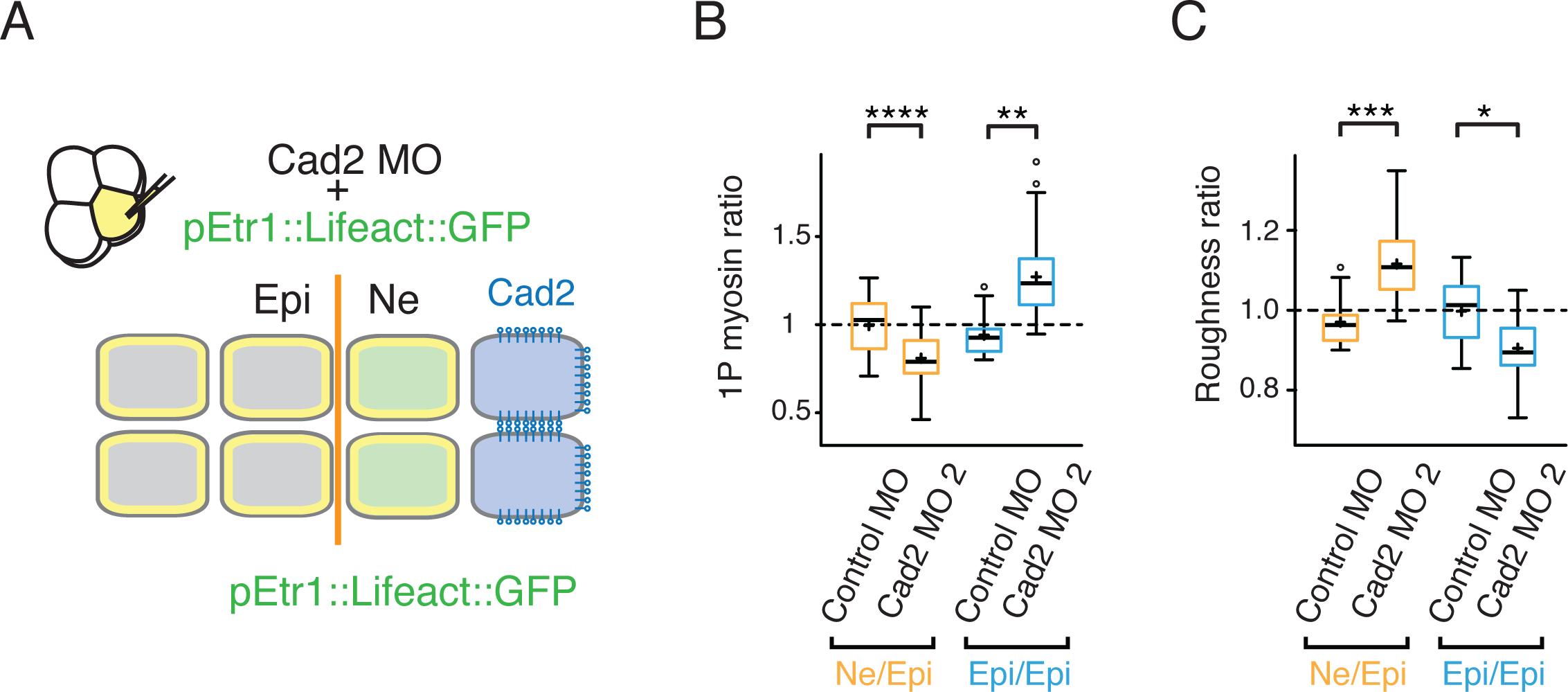
Controls for Cad2 MO specificity: (A) Co-injection of a second Morpholino Oligonucleotide (MO2) against Cad2 with an expression marker (pEtr1⸬Lifeact⸬GFP) into a single b4.2 blastomere at the 8-cell stage. (B, C) Box pots showing ratios of (B) 1P myosin and (C) boundary roughness on injected:non-injected sides of the embryo for the indicated junction or boundary types. Control measurements were made on embryos injected with control morpholino. (n > 23 junctions, from > 9 embryos for each condition in B; n >10 embryos for each condition in C. *p < 0.1, **p < 0.05, *** p < 0.005, **** p < 0.0005, Student’s t test.

**Figure S3 (related to Figure 5).**
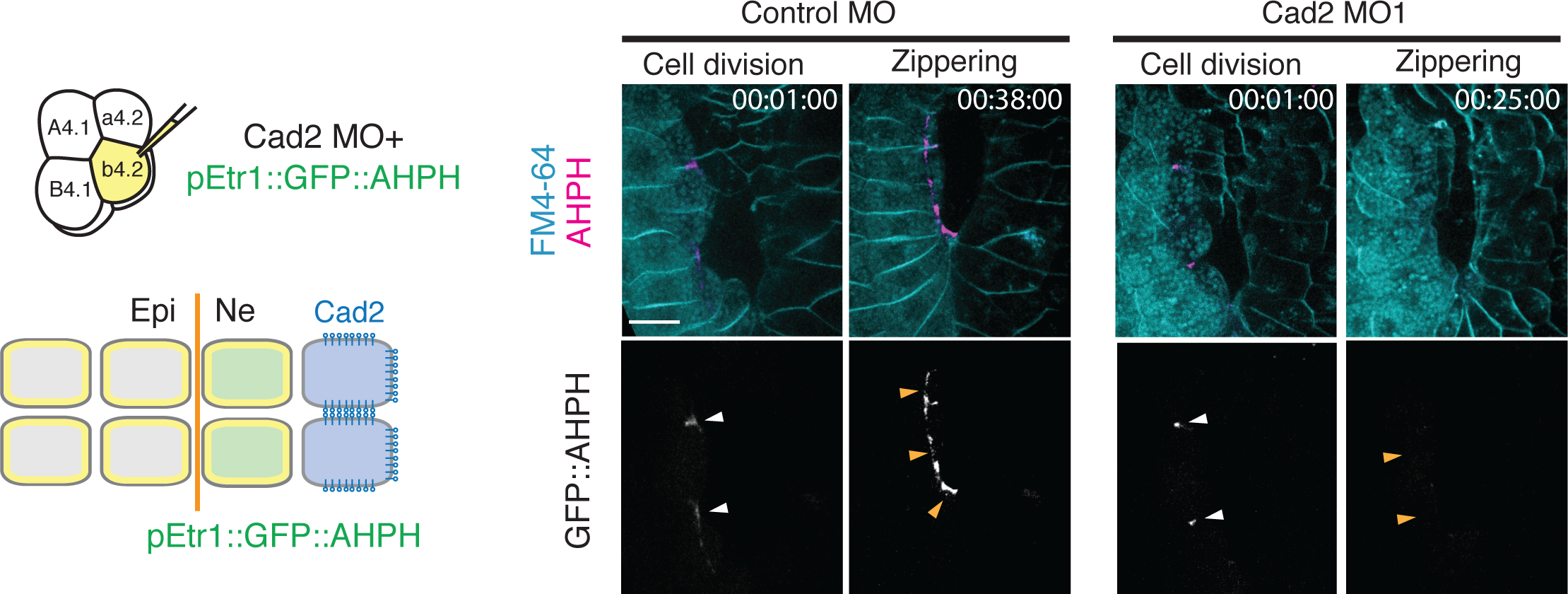
RhoA activity at cleavage furrow in Cad2 knockdown embryos. Co-injection of Morpholino Oligonucleotides (MO) against Cad2 with an expression marker (pEtr1⸬GFP⸬AHPH) into a single b4.2 blastomere at the 8-cell stage. Yellow indicates all descendants of the b4.2 blastomere; green indicates midline neural descendants that received the MO. Images extracted from Movie S6. Top row shows the biosensor (magenta) and a membrane marker (FM-464; blue); bottom row shows the biosensor alone. White and orange arrowheads indicate cleavage furrows and Ne/Epi junctions. Scale bars = 5 μm.

**Figure S4 (related to Figure 6).**
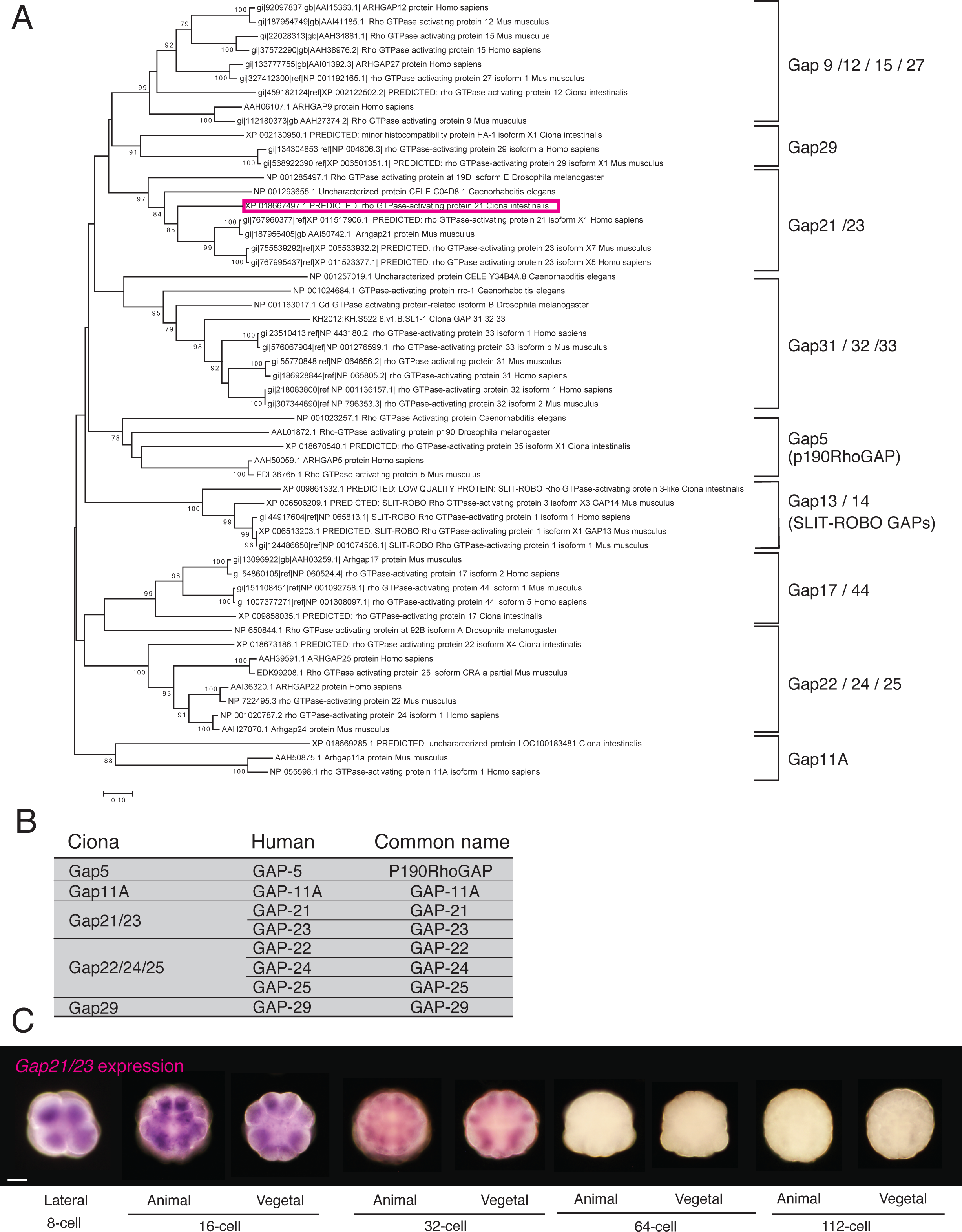
Phylogenetic analysis, expression pattern and localization of Gap21/23. (A) Sequence-based phylogeny for GAP domains from metazoan Rho GTPase activating proteins. Position of *Ciona robusta Gap21/23* is marked with a pink square. (B) List of *Ciona robusta* Rho GAPs tested in this study. (C) *Gap21/23* gene expression pattern from eight-cell to gastrula stage. Scale bars = 25 μm.

**Figure S5 (related to Figure 7).**
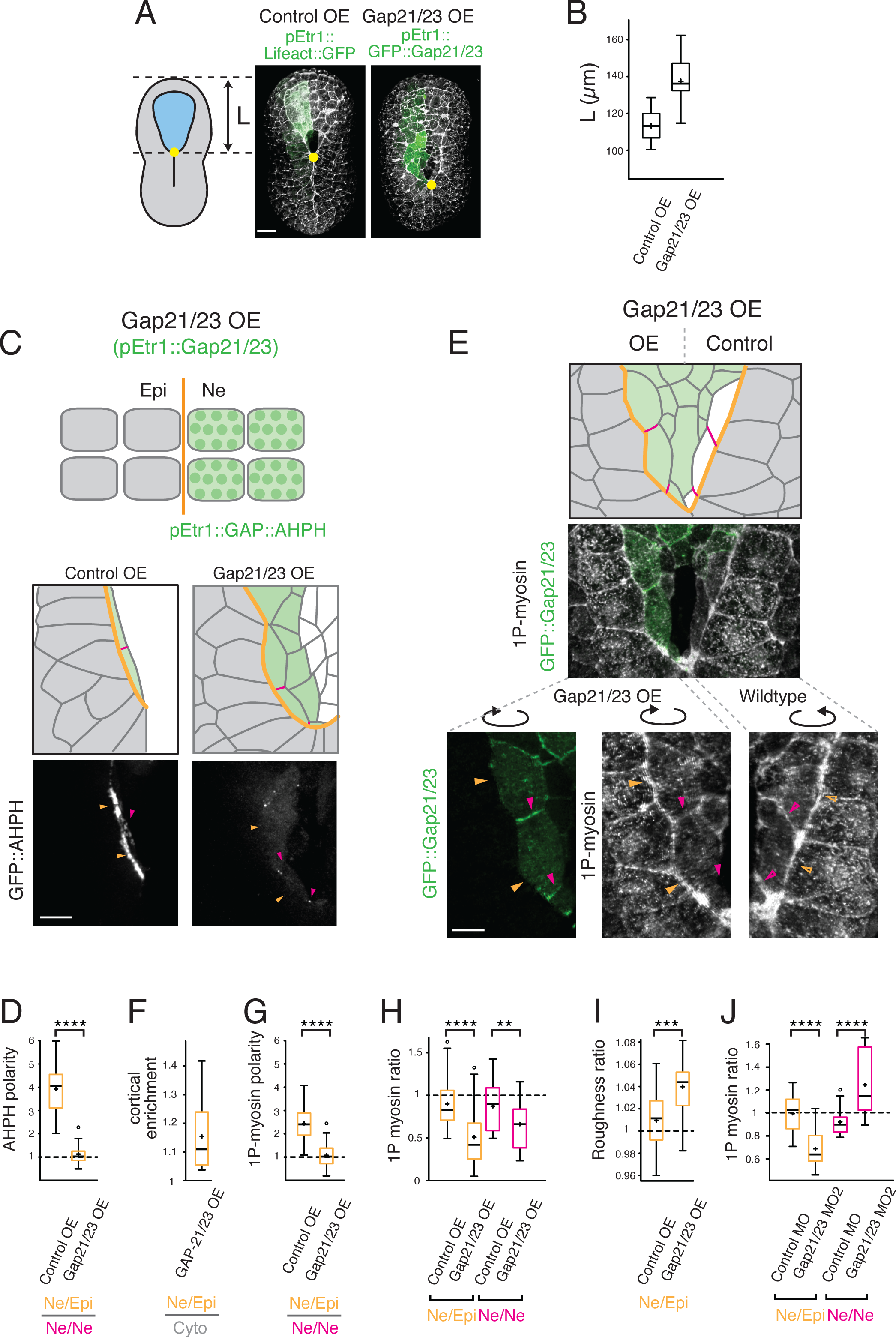
Gap21/23 over-expression in midline neural cells abolishes RhoA/myosin II polarity and blocks zippering. (A) GFP⸬Gap21/23 (green) is over-expressed in neural cells on one side of the embryo. Left: schematic showing measurement of “un-zippered length” L. Right: Embryos over-expressing pEtr1⸬GFP⸬Gap21/23 or a control transgene (pEtr1⸬Lifeact⸬GFP), then fixed and co-stained with phalloidin. Scale bars = 10 μm. (B) Quantitation of zipper progression in Gap21/23 over-expressing embryos (n > 14 embryos for each condition). (C, D) Effects of Gap21/23 over-expression on distribution of GFP⸬AHPH in midline neural cells. Top schematic: green indicates cells expressing pEtr1⸬GFP⸬AHPH and pEtr1⸬Gap21/23. Bottom panels: Orange lines and arrowheads indicate Ne/Epi junctions and magenta lines and arrowheads indicate Ne/Ne junctions in midline neural cells. Scale bars = 5 μm. (D) GFP⸬AHPH polarity measured as the ratio of intensities on Ne/Epi⸬Ne/Ne junctions. (E-H) Effects of Gap21/23 over-expression on distribution of 1P-myosin in midline neural cells. (E) Top panel shows dorsal surface view of a neurula stage embryo, over-expressing GFP⸬Gap21/23 in all neural cells except midline neural cells on one side of the embryo, then fixed and immunostained for 1P myosin. Green color fill indicates neural cells expressing GFP⸬Gap21/23. Orange line and arrowheads indicate the original Ne/Epi boundary, magenta lines and arrowheads indicate Ne/Ne junctions in midline neural cells. Second panel: superposition of GFP injection marker and 1P-myosin. Third panel shows localization of GFP⸬Gap21/23 (left) or 1P-myosin (right) in the same embryo. Open and filled arrowheads indicate corresponding boundaries on GFP⸬Gap21/23 over-expressing and control sides of the embryo. Scale bars = 5 μm. (F) Relative enrichment of GFP⸬Gap21/23, measured as the ratio of junctional to cytoplasmic intensity. (G) 1P myosin polarity measured as the ratio of intensities on Ne/Epi⸬Ne/Ne junctions. (H) 1P myosin ratio in Gap21/23-expressing embryos. (I) Roughness ratio in Gap21/23-expressing embryos. (n > 14 embryos for each condition) (J) 1P myosin ratio in embryos injected with a second Morpholino Oligo (MO2) against Gap21/23. (n > 19 junctions from > 8 embryos for each condition in D, F-J). **p < 0.05, **** p < 0.0005, Student’s t test.

**Figure S6 (related to Figure 7).**
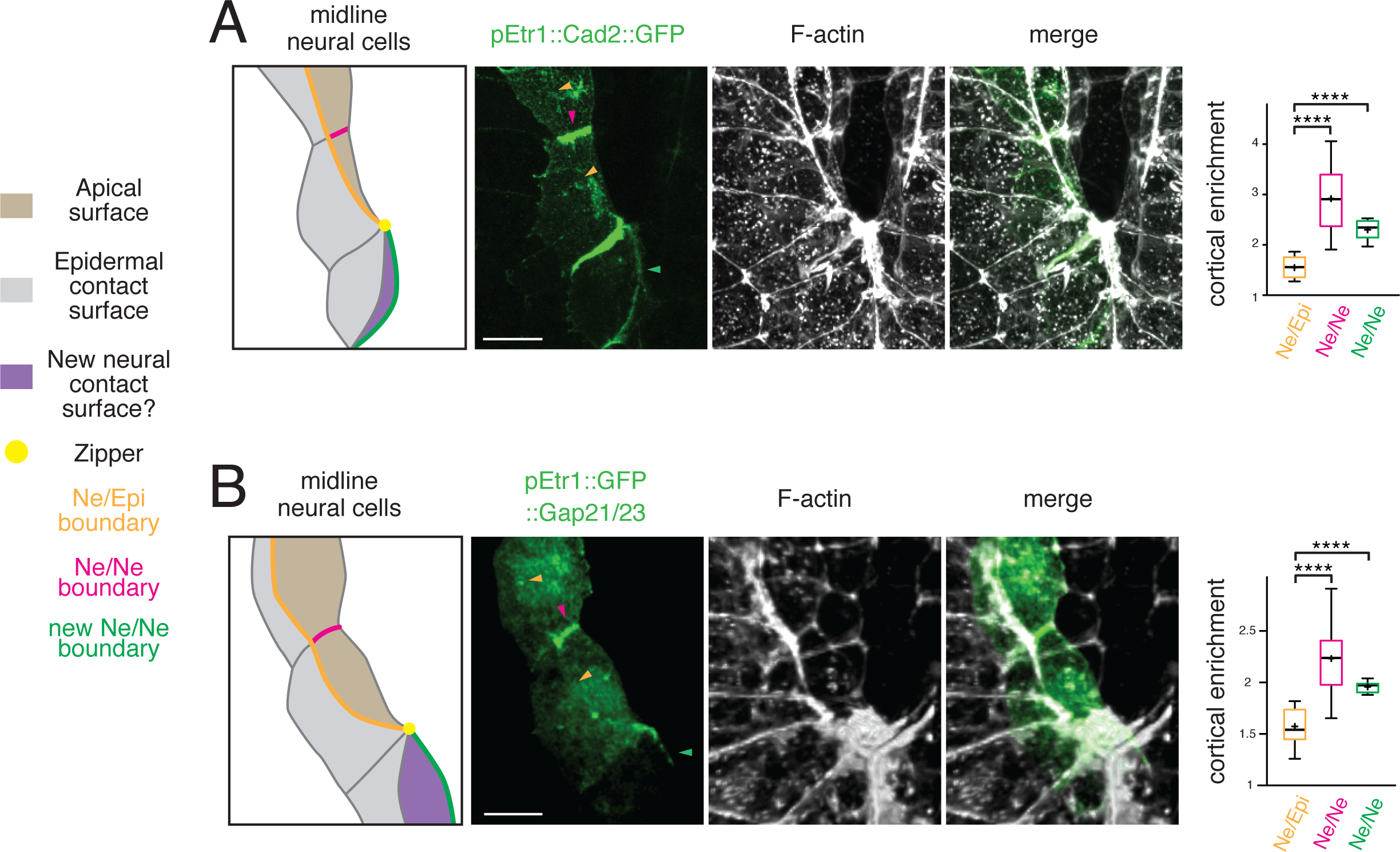
Dynamic recruitment of Cad2 and Gap21/23 to new Ne/Ne junctions behind the zipper. (A, B) Neurula stage embryos expressing pEtr1⸬Cad2⸬GFP (A) or pEtr1⸬GFP⸬Gap21/23 (B), fixed and counter-stained with phalloidin. Left panel: Manual tracing of midline neural cells on the left side of the embryo ahead of and behind the zipper. Color fill indicates apical surface (brown), Epidermal contact surfaces (grey) and newly-formed surfaces of contact with neural cells on the opposite (right) side just behind zipper (purple). Orange lines and arrowheads indicate the original Ne/Epi boundary. Magenta lines and arrowheads indicate Ne/Ne junctions ahead of zipper. Green lines and arrowheads indicate newly formed Ne/Ne junctions behind zipper. Yellow circle indicates zipper. Right three panels show Cad2⸬GFP (GAP⸬Gap21/23) signal, phallodin stain, and merge. Box plots show relative enrichment of Cad2⸬GFP and GAP⸬Gap21/23, measured as the ratio of junctional to cytoplasmic intensity for junction types indicated below each graph. (n > 12 junctions, from > 6 embryos in A; n >5 embryos in B). **** p < 0.0005, Student’s t test. Scale bars = 5 μm.

**Movie S1 (related to Figure 1).** Myosin II dynamics during zippering in live embryos expressing iMyo⸬GFP in epidermal cells on one side of the embryo (left) or in all midline neural cells (right). Each frame is the maximum intensity projection of 15 images collected at 0.75 μm intervals in Z near the apical surface; frames were collected at 60s intervals. The movie is displayed at 15 fps.

**Movie S2 (related to Figure 3).** 3D reconstruction of a neurula stage embryo in which Cad2 MO was injected into one b4.2 blastomere at the 8-cell stage. Orange arrowheads indicate Ne/Epi junctions on injected (filled orange arrowheads) and non-injected (open orange arrowheads) sides of the embryo. Blue arrowheads indicate junctions between injected and non-injected neural cells (filled blue arrowheads) and corresponding junctions on the non-injected control side (open blue arrowheads) sides of the embryo. See also Figure 3C.

**Movie S3 (related to Figure 3).** Time-lapse movies of embryos in which either a control (left) or Cad2 (right) morpholino was co-injected with pEtr1⸬Lifeact⸬GFP into a single b4.2 blastomere at the 8-cell stage. Embryos were counter-stained with FM-464 and imaged as described in Materials and Methods. GFP expression marks neural cells receiving the morpholino. Each frame is a maximum intensity projection of 15 images collected at 0.75 μm intervals in Z near the apical surface; frames were collected at 60s intervals. The movie is displayed at 15 fps.

**Movie S4 (related to Figure 4).** Time lapse movie of junctional dynamics during zippering in an embryo expressing ZO1⸬GFP under the control of a promoter (pFOG) that drives expression in all epidermal cells and a subset of neural cells lying along the Ne/Epi boundary. Because of mosaic transgene expression, only half of the embryo expresses ZO1⸬GFP. Each frame is the maximum intensity projection of 15 images collected at 0.75 μm intervals in Z near the apical surface; frames were collected at 30s intervals. The movie is displayed at 25 fps.

**Moive S5 (related to Figure 5).** Visualization of active RhoA dynamics during zippering in live embryos expressing GFP⸬AHPH under the control of a neural-specific promoter (pEtr1) in midline neural cells. Left panel shows GFP⸬AHPH (magenta) with a counter-stain FM-464 (cyan). Right panel shows GFP⸬AHPH alone. Each frame is the maximum intensity projection of 15 images collected at 0.75 μm intervals in Z near the apical surface; frames were collected at 60s intervals. The movie is displayed at 15 fps.

**Movie S6 (related to Figure 5).** Time-lapse movies showing active RhoA dynamics in embryos in which either a control (left) or Cad2 (right) morpholino was co-injected with pEtr1⸬GFP⸬AHPH into a single b4.2 blastomere at the 8-cell stage. Embryos were counter-stained with FM-464 and imaged as described in Materials and Methods. pEtr1⸬GFP⸬AHPH is expressed only in midline neural cells, on the left side of the embryo, that received the morpholino. Each frame is the maximum intensity projection of 15 images collected at 0.75 μm intervals in Z near the apical surface; frames were collected at 60s intervals. The movie is displayed at 15 fps.

**Movie S7 (related to Figure 7).** 3D reconstruction of a neurula stage embryo in which Gap21/23 MO was injected into one b4.2 blastomere at the 8-cell stage. Orange arrowheads indicate Ne/Epi junctions on injected (filled orange arrowheads) and non-injected (open orange arrowheads) sides of the embryo. Magenta arrowheads indicate junctions between Gap21/23 MO-injected and non-injected neural cells (filled magenta arrowheads) and corresponding junctions on the non-injected control side (open magenta arrowheads) sides of the embryo. See also Figure 7E.

